# DNA Methyltransferase Inhibition During CAR T-cell Manufacturing Enhances Antitumor Activity in Immunocompetent Brain and Ovarian Cancer Models

**DOI:** 10.64898/2026.11.23.768631

**Authors:** Elton VanNoy, Zhongzhen Yi, Abigail Lee, Nathaniel Barton, Kaleem Coleman, Lynne El Ghorayeb, Katherine B. Chiappinelli, Dalia Haydar

## Abstract

Limited persistence and progressive dysfunction remain major barriers to effective CAR T-cell therapy for solid tumors. Epigenetic regulation, particularly DNA methylation, can reinforce dysfunctional T-cell programs by restricting expression of genes required for sustained effector function. CAR T-cell manufacturing provides an opportunity to pharmacologically prime CAR T-cells prior to infusion without additional treatment burden. Therefore, we hypothesized that transient DNA methyltransferase (DNMT) inhibition during CAR T-cell manufacturing would epigenetically modulate CAR T-cell state and enhance functional and antitumor activity against solid tumors. We evaluated transient DNMT inhibition during *ex vivo* murine CAR T-cell manufacturing using two mechanistically distinct compounds: the nucleoside analogue 5-azacytidine (5-Aza) and the selective DNMT1 inhibitor GSK3685032 (GSK). Epigenetic and transcriptomic profiling, together with functional analyses including repeated antigen stimulation, were performed to characterize the molecular and functional effects of DNMT inhibition on CAR T-cells. Therapeutic efficacy was evaluated in immunocompetent syngeneic murine models of pediatric glioma and ovarian cancer by assessing tumor burden and survival. 5-Aza treatment induced dose-dependent epigenetic and transcriptional changes in CAR T-cells that were accompanied by enhanced expansion, sustained cytotoxicity and cytokine production during repeated antigen challenge *in vitro*. These improvements translated *in vivo*, with enhanced tumor control across immunocompetent models of high-grade glioma, diffuse midline glioma, and ovarian cancer using both B7-H3- and GD2-targeting CAR constructs. The dose associated with the greatest benefit varied by CAR construct and tumor model. In contrast, selective DNMT1 inhibition with GSK did not recapitulate the molecular or functional benefits of 5-Aza and reduced therapeutic efficacy *in vivo*. Collectively, these findings demonstrate that transient epigenetic modulation during CAR T-cell manufacturing can durably reshape CAR T-cell state and enhance antitumor efficacy. Broad DNMT inhibition with 5-Aza produced molecular, functional, and therapeutic benefits that were distinct from selective DNMT1 inhibition with GSK. These results support transient 5-Aza treatment during manufacturing as a translatable strategy to improve CAR T-cell therapies for solid tumors.

## INTRODUCTION

Chimeric antigen receptor (CAR) T-cell therapy has achieved durable clinical responses in hematologic malignancies, leading to multiple regulatory approvals^1,2^. However, translating these successes to solid tumors has remained challenging. CAR T-cell therapies targeting solid tumors have demonstrated encouraging safety profiles in clinical trials but have generally yielded modest and transient antitumor responses^3–6^. This underscores the need for clinically translatable strategies that enhance CAR T-cell efficacy without substantially increasing manufacturing complexity or patient treatment burden. This need is particularly acute in aggressive solid malignancies such as pediatric diffuse midline glioma (pDMG) and ovarian cancer. pDMG has a median survival of approximately 12 months, with radiotherapy remaining the only standard-of-care treatment^7^. In ovarian cancer, approximately 70% of patients experience disease recurrence following standard therapy, and treatment-resistant disease remains a major clinical challenge^8,9^.

CAR T-cell efficacy is frequently constrained by diminished function following repeated stimulation and limited persistence^10^. Epigenetic regulation plays a central role in T-cell differentiation, function, and persistence^11,12^. In particular, DNA methylation contributes to the establishment and maintenance of T- cell fate, with altered methylation programs contributing to dysfunctional states that restrict T-cell fitness^13–15^. The *ex vivo* activation and expansion required for CAR T-cell production can also induce progressive differentiation and epigenetic changes that may limit durability of the final product^16^. Because less differentiated T-cell states are associated with improved expansion, persistence, and antitumor activity following adoptive transfer^17,18^, modulation of the T-cell methylome represents a potential strategy for improving CAR T-cell therapy.

DNA methylation is added by DNA methyltransferases (DNMTs), with DNMT1 predominantly maintaining established methylation patterns and DNMT3A/B adding de novo methylation^19,20^. Genetic disruption of DNMT3A can preserve favorable T-cell states and enhance CAR T-cell persistence and antitumor activity^21^. Several small-molecule DNMT inhibitors (DNMTis) are FDA approved for hematologic malignancies^22^ and have been investigated as tumor-sensitizing agents to enhance immune-mediated killing^23,24^. However, fewer studies have examined DNMT inhibition as a strategy to directly reprogram T-cells. Notably, decitabine exposure during CD19 CAR T-cell manufacturing altered T-cell differentiation and enhanced antitumor activity in preclinical models of acute lymphoblastic leukemia and non-Hodgkin lymphoma^25^. These findings support pharmacologic methylome modulation during CAR T-cell manufacturing, but its effects across distinct CAR platforms and within solid tumor settings remain poorly defined.

The *ex vivo* manufacturing process provides a controlled setting to transiently modify CAR T-cells before infusion. 5-Azacytidine (5-Aza) is an FDA-approved nucleoside analog with broad DNA methylation-modulating activity^26^, whereas GSK3685032 (GSK) is an experimental non-nucleoside inhibitor selective for DNMT1^27^. Here, we evaluated transient DNMT inhibition during the manufacturing of murine CAR T-cells targeting B7-H3 and GD2, clinically relevant antigens expressed across pediatric and adult solid tumors^28,29^. The use of syngeneic, immunocompetent tumor models further enables evaluation of CAR T-cell function and therapeutic efficacy within the context of an intact immune system. By testing distinct DNMT inhibitors across CAR constructs and tumor models, we sought to determine whether transient methylome modulation can improve CAR T-cell function against solid tumors. This approach provides an opportunity to reprogram the CAR T-cell product before infusion, with the potential to enhance therapeutic function while limiting systemic drug exposure.

## METHODS

### Animal Models

All animal studies were conducted under protocols approved by the Institutional Animal Care and Use Committee (IACUC) at Children’s National Research Institute (Protocol #00030512) and The George Washington University (#A2024-11, #A2011-019). Animals were euthanized at predefined endpoints, including ≥20% body weight loss, neurological symptoms, or veterinary recommendation. C57BL/6J and B6(Cg)-Tyrc-2J/J (albino) mice (The Jackson Laboratory), 5–15 weeks of age and of both sexes, were used for syngeneic studies.

Briefly, for orthotopic glioma models, 1 × 10^5^ GL261 or KPAP.ffLuc tumor cells were stereotactically implanted into the cortex (HGG) or pons (DMG), as described previously^30^. GL261- and KPAP.ffLuc- bearing mice received 1-2 × 10^6^ CAR T-cells intratumorally 14 or 10 days after tumor implantation, respectively. For ovarian cancer studies, female C57BL/6J mice received 5 × 10^6^ ID8 *Trp53*^−/− cells intraperitoneally, followed 21 days later by 3 × 10^6^ CAR T-cells intraperitoneally. Therapeutic efficacy was primarily assessed by survival, with bioluminescence imaging used to monitor tumor burden in ffLuc-expressing models. Detailed procedures, animal monitoring and predefined humane endpoints are provided in the <u>Supplementary Materials and Methods.</u>

### Murine CAR T-cell Generation

Second-generation murine CARs were generated using retroviral vectors encoding an m276-derived (B7H3) or 14G2a-derived (GD2) scFv linked to CD28 costimulatory domains. Murine T-cells were selectively purified from mouse spleens using the Pan T-Cell Isolation Kit II (130-095-130, Miltenyi Biotec) as previously described^29^. Isolated T-cells were washed and activated using plate-bound CD3/CD28 antibodies (553057 and 553294 respectively, BD Biosciences) in complete RPMI media supplemented with 50 U/mL of human IL-2 (Ro 23-6019 NIH (NCI)). Following 48 hours of activation, T-cells were transduced with GPE-86-derived retroviral particles using RetroNectin-coated plates (T100B; Takara Bio). Beginning on the day of transduction, cells received 5-azacytidine (5-Aza; A2385, Sigma-Aldrich) or GSK3685032 (GSK; E1046, Selleck Chemicals) once daily for 3 consecutive days at concentrations of 50, 100 and 200nM. Forty-eight hours after transduction, cells were harvested, re- plated for expansion, and maintained in recombinant human IL-2 (50 U/mL) until Day 8 post- transduction; the final DNMTi dose was administered following replating. CAR expression was assessed by flow cytometry on days 5-6 post-transduction. Unless otherwise indicated, *in vitro* and *in vivo* studies were initiated on days 5-8 post-transduction. On day 6 post transduction, CAR T-cell pellets were collected for western blot analysis of DNMT inhibition from drug treatment. Drug preparation, western blotting and T-cell flow cytometry–based analyses are described in the <u>Supplementary</u> <u>Materials and Methods.</u>

### Functional Analysis of Murine CAR T-cells *In Vitro*

#### Serial Co-culture Assay

To perform repeated stimulation assays, 5e^5^ tumor cells (either GL261 or MBR525-1) were seeded in a 24-well plate and allowed to attach to the bottom of the well for 2-4 hours at 37°C. Cell line acquisition and culture conditions are detailed in the <u>Supplementary Materials and Methods.</u> CAR T-cells were then washed with cytokine-free media, and 1e^6^ T-cells were introduced into the wells containing the tumor cells. Repeated stimulations were performed by re-plating fresh tumor cells and re-establishing co-culture with collected T-cells at the same E:T ratio every 2-4 days. Stimulations continued until T- cell proliferation and/or killing activity were no longer observed. In instances where fewer than 1e^6^ T- cells were collected but proliferation and/or killing activity persisted, all remaining cells were re- suspended in 1mL and plated on tumor cells at the maximum ratio achievable.

#### Cytotoxicity Assay

Colorimetric MTS reagent [3-(4,5-dimethylthiazol-2-yl)-5-(3-carboxymethoxyphenyl)-2-(4-sulfophenyl)- 2H-tetrazolium] (Promega, G3581) was used to quantify the cytotoxicity of CAR T-cells. Briefly, GL261 or MBR525-1 tumor cells were added to a 96-well tissue culture-treated plate and allowed to adhere for 2-4 hours. T-cell cultures post-manufacturing or post-stimulation were then added to the tumor cells at varying E:T ratios (2:1, 1:1, 0.5:1, 0.25:1, and 0.125:1, performed by serial dilution). T-cells and tumor cells were incubated for 72 hours at 37°C. After incubation, T-cells were removed by gently pipetting up and down to avoid disrupting remaining adherent tumor cells. MTS reagent was then diluted in complete media and added to each well, incubated at 37°C for 2h and absorbance was then measured at 492nm by using a SPARK Plate Reader (Tecan). The percentage of live tumor cells was calculated using the following formula: [(Absorbance of sample- Absorbance of media only)/ (Absorbance of tumor only - Absorbance of media only)] x100.

#### Cytokine Production

During each cycle of the serial co-culture assay, supernatant was gathered 24 hours after coculturing T-cells with tumor cells and frozen at −20°C. The evaluation of IL2, TNFa and IFNy cytokine production was carried out using mouse Quantikine ELISA kits (R&D Systems; SM2000, SMTA00Ba, SMIF00, respectively). Optical density was measured at 540nm by using a SPARK Plate Reader (Tecan) and cytokine concentrations were calculated from standard curves generated for each assay.

### Statistics

Statistical analyses were performed using GraphPad Prism (GraphPad Software, San Diego, CA, USA). Data are presented as mean ± SEM unless otherwise indicated. Comparisons between two groups were performed using paired or unpaired two-tailed tests, as appropriate. For experiments involving three or more groups, ordinary or repeated-measures one-way ANOVA with multiple- comparison correction was used for normally distributed data. Donor-matched datasets normalized to paired controls were analyzed using Wilcoxon signed-rank tests or one-sample tests against a theoretical value of 1, as appropriate. For experiments involving two independent variables, two-way ANOVA or mixed-effects models with multiple-comparison correction were performed as indicated in the figure legends. Normality was assessed using GraphPad Prism, and appropriate nonparametric tests were used when data did not meet assumptions of normality. Survival was analyzed using Kaplan-

Meier curves and compared using the log-rank (Mantel-Cox) test. A two-sided *P* value < 0.05 was considered statistically significant. Significance is denoted as follows: *P* < 0.05 (\**), P < 0.01 (**), P < 0.001 (\*\*\**), *P* < 0.0001 (****), and ns (not significant). Sequencing analyses workflows are described in detail in the <u>Supplementary Materials and Methods.</u>

## RESULTS

### DNMT inhibition during CAR T-cell manufacturing does not negatively impact T-cell transduction or proliferation

Murine T-cells were activated for 48 hours followed by transduction of a second-generation CD28.ζ CAR with an m276 (B7H3) antigen recognition domain, as previously described described^29^. To incorporate DNMT inhibition without compromising CAR T-cell activation, 5- Aza was administered during the 3-day transduction phase (Figure 1A). Dose-titration studies demonstrated that 200nM 5-Aza was the maximally tolerated concentration before cytotoxic effects emerged (Supplementary Figure 1); therefore, 0, 50, 100, and 200nM were selected for subsequent experiments. B7H3 CAR transduction remained consistent across treatment groups, with a mean transduction efficiency of 58% (SD ±13.43%, Figure 1B). To confirm sustained DNMT inhibition following 5-Aza exposure, DNMT1 and DNMT3A protein abundance was evaluated by western blot 3 days after treatment cessation (day 6 post-transduction). Both proteins showed progressive reductions with increasing 5-Aza concentration (Figure 1C-D). Relative to untreated controls, DNMT1 abundance was significantly reduced at 200nM (*p* = 0.0048), whereas DNMT3A abundance was significantly reduced at all concentrations; 50nM (*p* = 0.0099), 100nM (*p* = 0.0032), and 200nM (*p* = 0.0003). Viability was not substantially altered (Figure 1E), and expansion was maintained at 50 and 100nM but reduced at 200nM relative to untreated controls (p=0.0273). Accordingly, 200nM was retained for *in vitro* studies but excluded from *in vivo* evaluation. 5-Aza treatment did not substantially alter the baseline phenotypic composition of the CAR T-cell product; however, modest elevations in exhaustion markers (TIM3 and PD1) were observed with increasing dosage (Supplementary Figure 2). To assess whether the effects of 5-Aza were independent of antigen receptor signaling, we performed parallel manufacturing and characterization studies using a B7H3 Control CAR (Ctrl-CAR) containing a truncated endodomain. 5-Aza treatment did not significantly affect Ctrl-CAR transduction efficiency, cell viability, or expansion during *ex vivo* culture, consistent with the findings observed with the functional B7H3 CAR construct (Supplementary Figure 3A-C).

**Figure 1.**
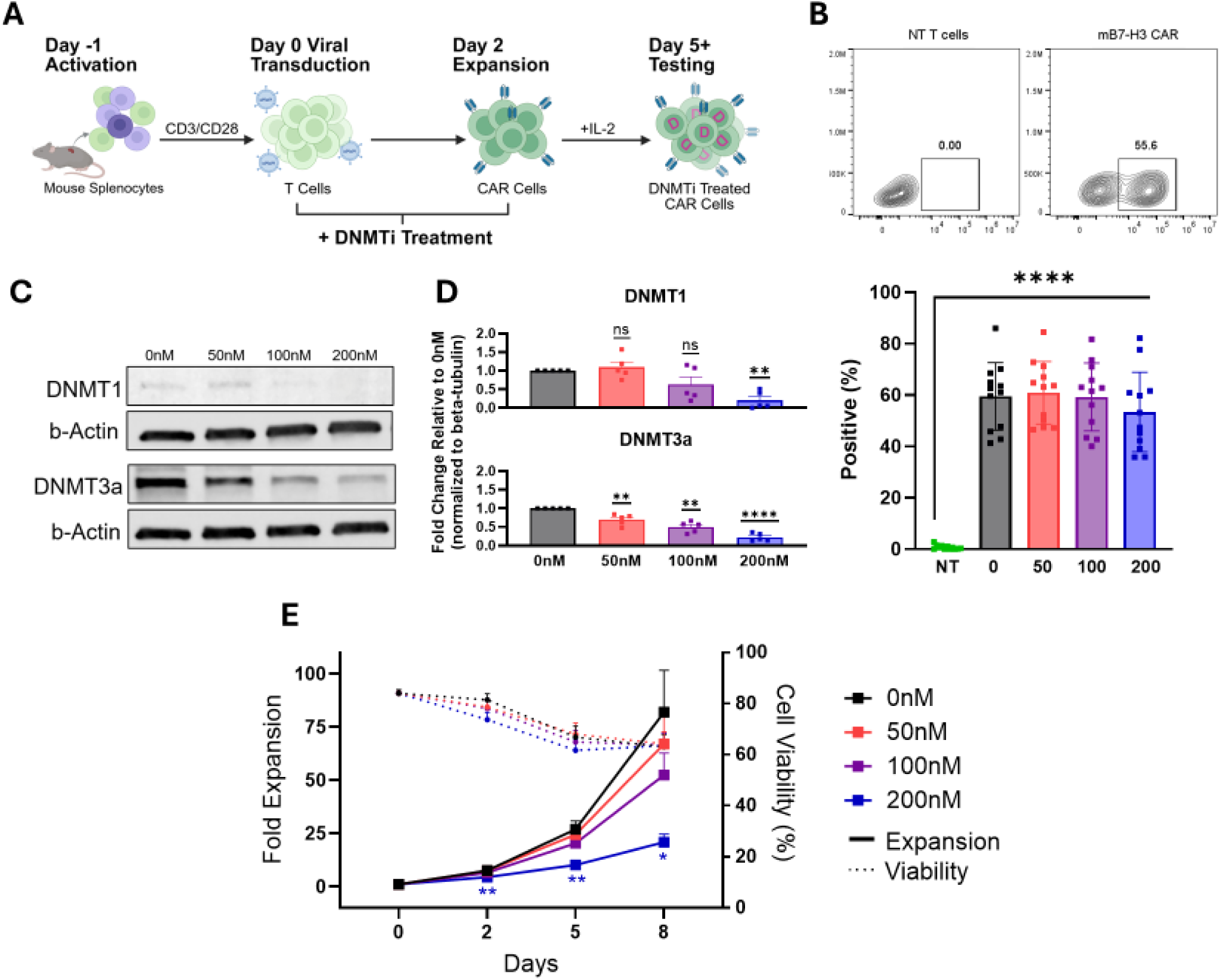
Characterization of 5-Aza treated mB7H3 CAR T cells. A) Schematic of DNMTi-treated CAR T cell manufacturing timeline. 5-Azacytidine (5-Aza) was added beginning on Day 0, concurrent with CAR transduction, and administered daily on days 0-2. B) CAR expression evaluated using flow cytometry at days 5-6 post transduction. Representative flow plots for non-transduced (NT) and B7H3 CAR expression in murine T cells. Summary plot of %F(ab’)_2_-positive T cells (n = 12, mean ± SEM, 1-way ANOVA with Tukey’s test for multiple comparisons, ****P < 0.0001. C-D) Protein was extracted from CAR T-cells 6 days post transduction and immunoblots were run for DNMT1 and DNMT3a with α-tubulin as a loading control. C) Representative cropped blots are shown here, and black lines indicate where one part of the blot ends and another begins. D) DNMT1 and DNMT3A expression was normalized to the corresponding untreated (0nM) control. Treatment conditions were compared with the theoretical value of 1 using two-sided one-sample t-tests with Holm correction for multiple comparisons, (n = 5, mean ± SEM). E) CAR T cells were expanded with IL-2 (50 U/mL) for 8 days following initiation of transduction. Fold expansion and viability were monitored throughout culture (n = 12 donors, mean ± SEM). Expansion kinetics were analyzed by two-way repeated-measures ANOVA with Dunnett’s multiple-comparisons test comparing each treatment group to the 0nM control at each time point, *P < 0.05, **P < 0.01.

### 5-Aza induces dose-dependent transcriptional and epigenetic remodeling in B7H3 CAR T- cells

To characterize the transcriptional and epigenetic effects of 5-Aza during CAR T-cell manufacturing, B7H3 CAR T-cells treated with increasing 5-Aza concentrations (0-200nM) were profiled by bulk RNA-seq and Infinium Mouse Methylation BeadChip analysis. RNA-seq identified 111 genes with significant dose-dependent expression changes, including 92 genes that increased and 19 genes that decreased with escalating 5-Aza exposure (Figure 2A). In parallel, methylation profiling identified 507 differentially methylated regions (DMRs), all of which exhibited reduced methylation with increasing 5-Aza dose (Figure 2B). Despite these widespread transcriptional and epigenetic changes, overlap between individual differentially expressed genes and DMR-associated genes was limited, with only four genes shared between analyses (OR = 1.42, *p* = 0.32; Figure 2C). In contrast, pathway-level analyses demonstrated substantial concordance between datasets. Among 4,166 GO biological process terms tested, 26 pathways were significantly enriched in both transcriptomic and methylation analyses (OR = 7.3, *p* = 9.6e-11, Figure 2D). Shared pathways clustered into functional programs associated with leukocyte differentiation, immune effector activation, lymphocyte activation and proliferation, leukocyte cell-cell adhesion, and leukocyte proliferation (Figure 2E). To assess whether 5-Aza induced remodeling extended specifically to genes involved in CAR T-cell effector function, we examined a curated panel of effector function associated genes. While, this panel did not meet significance thresholds for differential expression or methylation, all 10 genes showed inverse expression-methylation trends across doses, including *Ifng* (r=-0.86, p=0.14), *Prf1* (r=-0.94, p=0.064), *Nkg7* (r=-0.86, p=0.14), and *Tbx21* (r=-0.90, p=0.097; Figure 2F).

**Figure 2.**
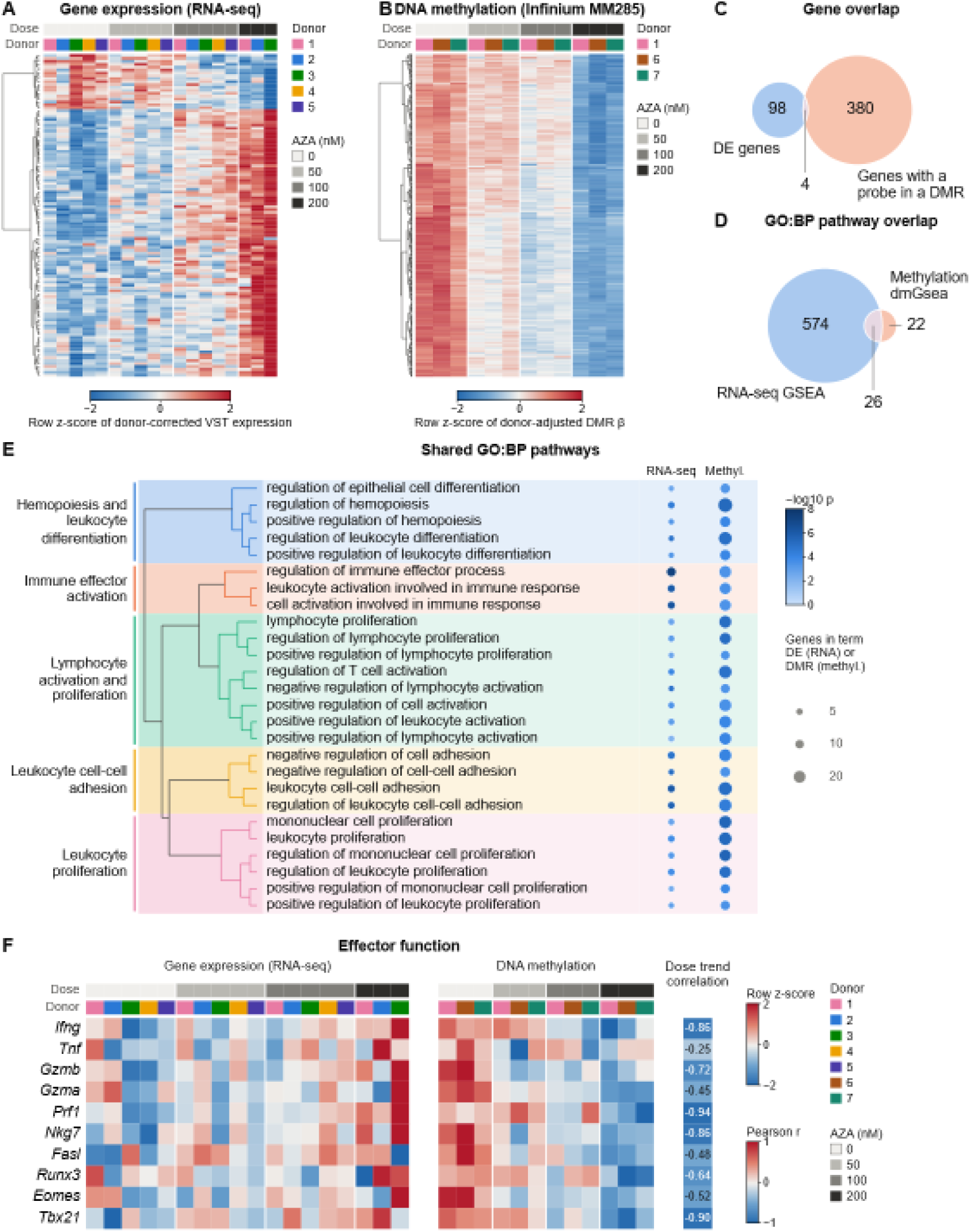
5-Azacitidine induces dose-dependent transcriptional activation and DNA hypomethylation of immune- activation pathways in B7H3 CAR T-cells. Mouse B7H3 CAR T-cells were treated with 5-azacytidine (AZA; 0, 50, 100, or 200nM) and profiled by bulk RNA-seq or with the Infinium Mouse Methylation BeadChip (MM285). Donors are numbered across both cohorts; donor 1 is the only shared donor. **A)** Heatmap showing the expression of 111 differentially expressed (DE) genes with AZA dose (92 up, 19 down; DESeq2, with dose as a continuous covariate). DE genes are defined as adjusted p < 0.05 and | log2FC/100nM AZA | > 0.5. Values are variance-stabilized expression with donor effect removed, z-scored per gene. Rows are clustered (Ward); columns are ordered by dose and annotated by donor. **B)** Heatmap showing methylation of 507 differentially methylated regions (DMRs). All DMRs are hypomethylated, consistent with AZA dose (DMRcate FDR < 0.05, | mean Δβ, 0→200 nM| > 0.10. n=12 samples (3 donors x 4 doses). Values are donor-corrected β values averaged across each DMR’s probes, z-scored per DMR. Rows are clustered (Ward); columns are ordered by dose and annotated by donor. **C)** Overlap between DE genes and genes with at least one promoter (1.5kb upstream of TSS) or gene-body probe inside a DMR (B), counted among the 13,699 genes measured on both platforms (102 of 111 DE genes; 384 of 441 DMR genes). One-sided Fisher’s exact test: OR = 1.42, p = 0.32). **D)** Overlap of GO biological-process (GO:BP) terms enriched (FDR < 0.05) in the DNA-seq (GSEA of genes ranked by DESeq2 dose statistic) and methylation (dmGsea, probes in the 507 DMRs against all gene-mapped probes, corrected for probe- number bias) analyses, among the 4,166 terms tested in both. One-sided Fisher’s exact test: OR = 7.3, p = 9.6e-11). **E)** The 26 shared GO:BP terms from (D). All are enriched among genes upregulated with dose (NES > 0) and among hypomethylated DMRs. Left, hierarchical clustering (Ward) of terms by GO semantic dissimilarity (1 - Wang similarity); colored blocks mark the five clusters, named by their dominant term keywords. Right, dot colors show -log10 p (RNA-seq GSEA; methylation dmGsea); dot sizes show the number of genes in the GO:BP term that are DE (RNA-seq) or have a promoter/gene-body probe in a DMR (methylation). **F)** Gene expression (left) and methylation (right) of a curated list of effector-function genes. Expression: donor-corrected log2-normalized counts (18 samples); methylation: donor-corrected median β over each gene’s promoter and gene-body probes (12 samples); each block z-scored per gene. Pearson r (right column) compares each gene’s mean RNA and methylation values across the four doses (n = 4 points per gene). Because the cohorts share only donor 1, r measures gene-level dose-response concordance rather than a paired-sample correlation. None of the genes are DE or have a probe in a DMR.

### 5-Aza priming enhances the persistence of B7H3 CAR T-cells against HGG *in vitro*

B7H3 CAR T-cells manufactured with 5-Aza were serially co-cultured with murine HGG cell line, GL261, at an effector to target (E:T) ratio of 2:1 without exogenous cytokines. GL261 was used as a model due to its reliable growth and adherence *in vitro.* However, all results found utilizing GL261 were validated in other tumor models *in vitro* and *in vivo*. Target antigen expression was confirmed on all utilized tumor cell lines by flow cytometry, and the corresponding CAR constructs are shown in Supplementary Figure 4. Ctrl-CAR T-cells, regardless of 5-Aza dosage, were unable to complete 1 repeat stimulation (Supplementary Figure 3D-E). B7H3 CAR T-cells generated with 50nM, and 100nM 5-Aza demonstrated more sustained proliferative fitness, with multiple donors maintaining cytotoxicity and expansion through ≥12 repeat stimulations and achieving greater peak fold expansion (Figure 3A). Consistent with the reduced *ex vivo* expansion observed during manufacturing, CAR T-cells treated with 200nM 5-Aza exhibited diminished functional persistence and supported fewer sequential GL261 tumor-cell stimulations. At the second stimulation, CAR T-cells manufactured with 50nM and 100nM 5-Aza demonstrated enhanced proliferation compared to untreated controls, achieving mean fold expansions of 7.3e6 and 6.7e6 cells, respectively, compared with 5.8e6 cells in the 0nM treatment group (*p* = 0.0273 and *p* = 0.0421, respectively; Figure 3B). This proliferative advantage became more pronounced with repeated antigen challenge, with 100nM-treated CAR T-cells reaching a mean fold expansion of 104e6 cells by the sixth stimulation compared with 27e6 cells in untreated controls (*p* = 0.0192; Figure 3B). Analysis of the maximum fold expansion achieved across all stimulations demonstrated that 50nM and 100nM 5-Aza significantly enhanced proliferative capacity relative to untreated controls (*p* = 0.0181 and *p* = 0.0203, respectively), whereas 200nM 5-Aza did not (p=0.0785; Figure 3C). The 50 and 100nM groups completed a mean of 5.6 stimulations, compared with 5.0 for untreated and 4.0 for 200nM-treated cells (Figure 3D). Phenotypic profiling following antigen stimulation revealed no substantial differences in memory phenotype or exhaustion marker expression across treatment groups (Supplementary Figure 5). 5-Aza-treated B7H3 CAR T-cells were also evaluated against the aggressive glioblastoma model, MBR525-1^31^ (Supplementary Figure 6).

**Figure 3.**
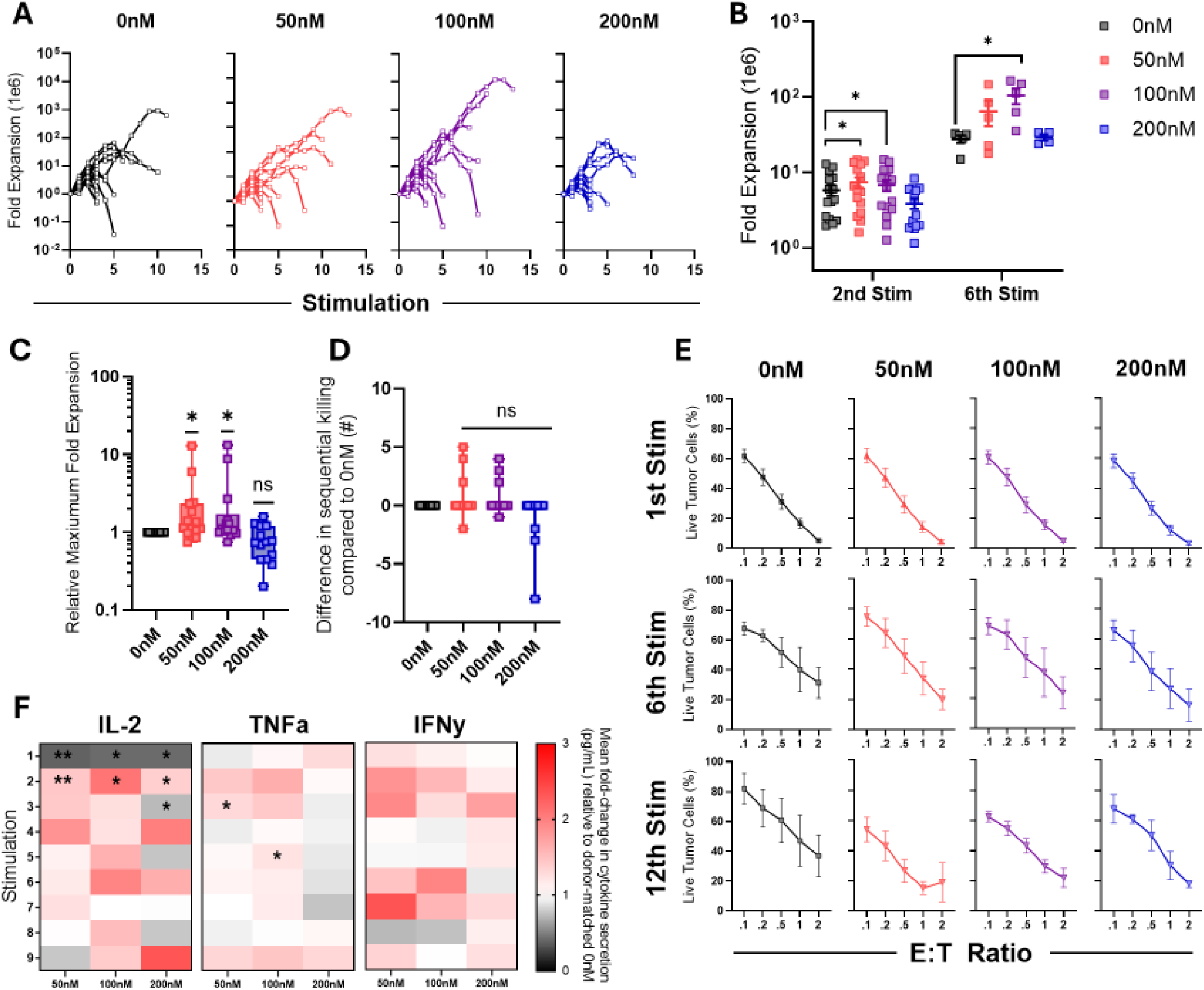
Functional characterization of 5-Aza treated B7H3 CAR T cells. Following 5-Aza treatment, B7H3 CAR T cells were serially challenged with GL261 tumor cells at a 2:1 effector-to-target ratio and restimulated every 2-3 days with new tumor cells until proliferative expansion and/or cytotoxic activity could no longer be sustained. **A)** Fold expansion across successive stimulation rounds is shown for individual donors (n = 10). **B)** Fold expansion at completion of the second and sixth stimulation rounds. Points represent individual donors (n = 10, mean ± SEM, one-way ANOVA with Dunnett’s multiple-comparisons test versus untreated controls (0nM)). Analyses of right-skewed datasets were conducted using a lognormal distribution model, *P < 0.05. **C)** Maximum fold expansion at each 5-Aza concentration relative to the donor-matched untreated control, calculated as the treated maximum divided by the non-treated maximum (n = 10, minimum-to-maximum range, one-sample Wilcoxon signed-rank test versus a hypothetical value of 1.0, *P < 0.05). **D)** Difference in the maximum number of sequential GL261 tumor-cell stimulations supported while maintaining CAR T-cell cytotoxic activity relative to non-treated (n = 15, minimum-to-maximum range, one-sample Wilcoxon signed-rank test versus a hypothetical value of 0). **E)** MTS cytotoxicity assay against GL261 tumor cells across indicated effector-to-target (E:T) ratios at the 1^st^, 6^th^ and 12^th^ stimulation (mean ± SEM, 1st stimulation, n = 17; 6th stimulation, n = 5; 12^th^ stimulation n = 3). **F)** IL-2, TNFα, and IFNγ secretion measured by ELISA over consecutive stimulations. Heatmap values represent the mean fold change in cytokine secretion for each treatment condition relative to the donor-matched untreated control at the corresponding stimulation (n = 2-5; one-sample t test versus a hypothetical value of 1.0, *P < 0.05).

Against this model, CAR T-cells manufactured with 50nM, and 100nM 5-Aza continued to achieve higher average maximum fold expansions (182e6 and 219e6 cells, respectively) than untreated controls (148e6 cells).

### 5-Aza treated CAR T-cells maintain cytotoxicity and alter cytokine secretion during repeated stimulation

To assess cytotoxicity and repeated tumor-killing capacity following chronic antigen stimulation, 72-hour MTS assays were performed at selected serial stimulations. All B7H3 CAR T-cell groups demonstrated efficient cytolytic activity against GL261 cells during initial stimulation, whereas control CAR T-cells did not (Supplementary Figure 3F). Repeated tumor exposure uncovered differences in functional persistence (Figure 3E). CAR T-cells generated with 50nM 5-Aza progressively separated from untreated controls and maintained superior tumor control through the 12th stimulation, particularly at challenging E:T ratios. In contrast, benefits observed with 100 and 200nM 5-Aza were not sustained, with cytolytic activity converging toward untreated levels after prolonged antigen challenge. Longitudinal cytokine secretion was assessed 24 hours after each stimulation. Following the first stimulation, IL-2 secretion was approximately 60% lower in all 5-Aza groups than untreated cells (foldchange, 0.38–0.42; all *p*<0.02). Following the second stimulation, IL- 2 increased 1.4-, 2.0-, and 1.4-fold in the 50, 100, and 200nM groups, respectively (all *p*<0.05; Figure 3F). 5-Aza-treated CAR T-cells, particularly the 50 and 100nM groups, also showed increases in TNFα and IFNγ across repeated stimulations. To explore the altered cytokine profile, we assessed cytokine gene methylation and expression (Supplementary Figure 7). Increasing 5-Aza doses were associated with reduced methylation at the *IL2* (*p*=0.007), *Ifng* (*p*=0.004), *Ifnb1* (*p*<0.001), and *Ifna* (*p*=0.03) loci. These changes resulted in modest transcriptional effects, with increased unstimulated *Ifng* expression (*p*=0.009), whereas cytokine transcript levels following antigen stimulation were not significantly altered.

### B7H3 5-Aza-treated CAR T-cells improve outcomes in immunocompetent brain tumor models

To evaluate the antitumor activity of 5-Aza-treated B7H3 CAR T-cells *in vivo*, we first used the syngeneic orthotopic GL261 model. Mice received 2e^6^ intratumoral B7H3 CAR T-cells 14 days after implantation (Figure 4A). Mice receiving untreated (0nM) CAR T-cells achieved a median survival of 39 days. More than 50% of animals in the 50nM and 100nM 5-Aza-treated groups remained alive at study completion, and median survival was therefore not reached (Figure 4B). Despite this improvement, survival did not significantly differ between either 5-Aza-treated group and non-treated CAR T-cells (*p*=0.6685). However, all CAR T-cell groups prolonged survival relative to tumor-only controls, with the strongest statistical evidence observed for the 100nM group (0nM, *p*=0.0038; 50nM, *p*=0.0091; 100nM, *p*=0.0002; Supplemental Figure 3G). No treatment-related toxicity was observed based on longitudinal body weight measurements (Figure 4C). Control CAR T-cells did not improve survival, regardless of 5-Aza exposure (median survival, 28–29 days; all *p*>0.05; Supplementary Figure 3G).

**Figure 4.**
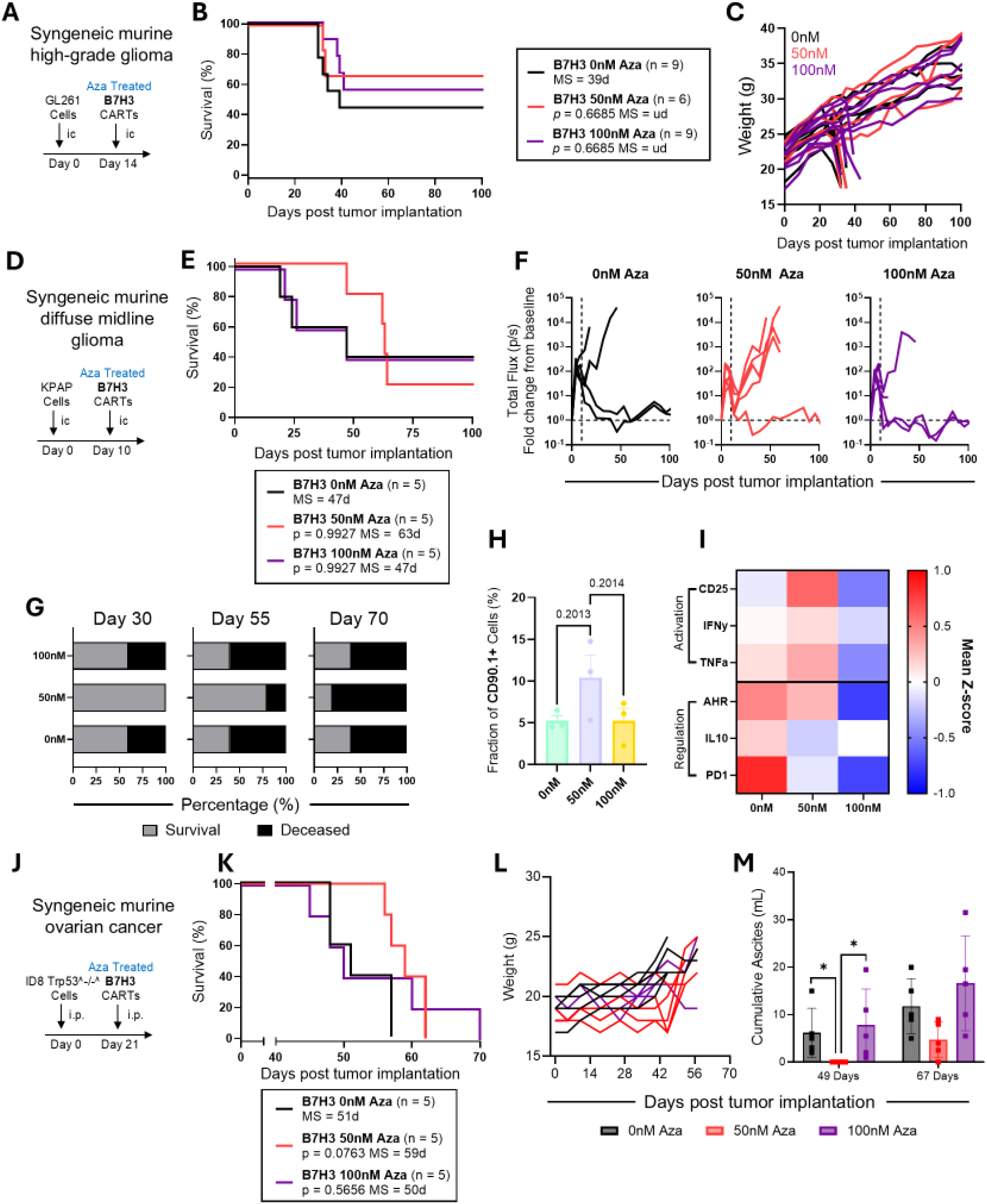
Evaluation of 5-Aza treated B7H3 CAR T cells *in vivo*. **A-C)** 1e5 GL261 cells were injected orthotopically into C57BL/6 mice, followed by intra-tumoral injection of 5-Aza treated B7H3 CAR T-cells. **A)** Experimental scheme. **B)** Kaplan-Meier survival curve (n = 6-9, log-rank Mantel-Cox test with Holm-Šídák for all pairwise multiple comparisons to 0nM, (MS = medial survival, ud = undefined). Three mice in the 50nM were excluded following unexpected early mortality that preceded mortality in the tumor-only control (Supplementary Figure 10) and occurred outside the anticipated tumor- related endpoint window. **C)** Changes in mice body weight determined weekly. **D-I)** 1e5 KPAP.ffLuc cells were injected orthotopically into Albino C57BL/6 mice, followed by intra-tumoral injection of 2e6 5-Aza treated B7H3 CAR T-cells. **D)** Experimental scheme. **E)** Kaplan-Meier survival curve (n = 5, log-rank Mantel-Cox test with Holm-Šídák correction for multiple comparisons to 0nM). **F)** Quantification of tumor burden over time baseline imaging. **G)** Summary plots showing percentage of surviving and deceased mice within each treatment group at days 30, 55 and 70 after tumor implantation. **H-I)** On day 4 following intracranial CAR T-cell administration, brains were harvested and dissociated for isolation of CD45+ cells, followed by comprehensive flow cytometric characterization. **H)** Quantification of intratumoral CAR T-cell infiltration, defined as CD90.1+ cells, across treatment groups (n=3, one-way ANOVA with Tukey test for multiple comparisons). **I)** Heatmap depicting z-score-normalized median fluorescence intensity (MFI) of phenotypic markers on infiltrating CD90.1+ CAR T cells. **J-M)** 5e6 ID8 *Trp53-/-* murine OC cells were injected intraperitoneally into female C57BL/6 mice, followed by intraperitoneal injection of 5-Aza treated B7H3 CAR T-cells. **J)** Experimental scheme. **K)** Kaplan-Meier survival curve (n =5, log-rank Mantel-Cox test with Holm-Šídák correction for multiple comparisons to 0nM). **L)** Changes in mice body weight determined weekly. **M)** Cumulative volume of ascites (fluid buildup in peritoneum) for treated mice at days 49 and 67 (n=5, repeated-measures one-way Kurskal-Wallis with Dunn’s multiple-comparisons test, *P < 0.05).

To evaluate 5-Aza-treated CAR T-cells in a pediatric DMG model, we next used the KPAP (H3.3^K27M^, <u>p</u>53^LOF^, <u>A</u>TRX^LOF^, <u>P</u>DGFRA^WT^) DMG model^32^. Given the neurosphere growth characteristics of KPAP cells, serial Annexin V assays demonstrated efficient B7H3-specific CAR T-cell cytotoxicity *in vitro* (Supplementary Figure 8). *In vivo,* KPAP tumors were implanted within the pons and longitudinally monitored by bioluminescence imaging and body weight following intratumoral administration of 2e^6^ CAR T-cells on day 10 (Figure 4D–F; Supplementary Figure 9). Mice receiving non-treated (0nM) or 100nM 5-Aza-treated CAR T-cells had a median survival of 47 days, whereas mice receiving 50nM 5- Aza-treated CAR T-cells had a numerically longer median survival of 63 days. However, survival did not significantly differ between either 5-Aza-treated group and the non-treated group (both p=0.9927; Figure 4E). However, mice receiving 50nM 5-Aza-treated CAR T-cells also showed delayed tumor burden progression (Figure 5F-G) and were the only CAR T-cell treatment group to significantly prolong survival relative to tumor-only controls (p=0.0062; Supplementary Figure 10A).

**Figure 5.**
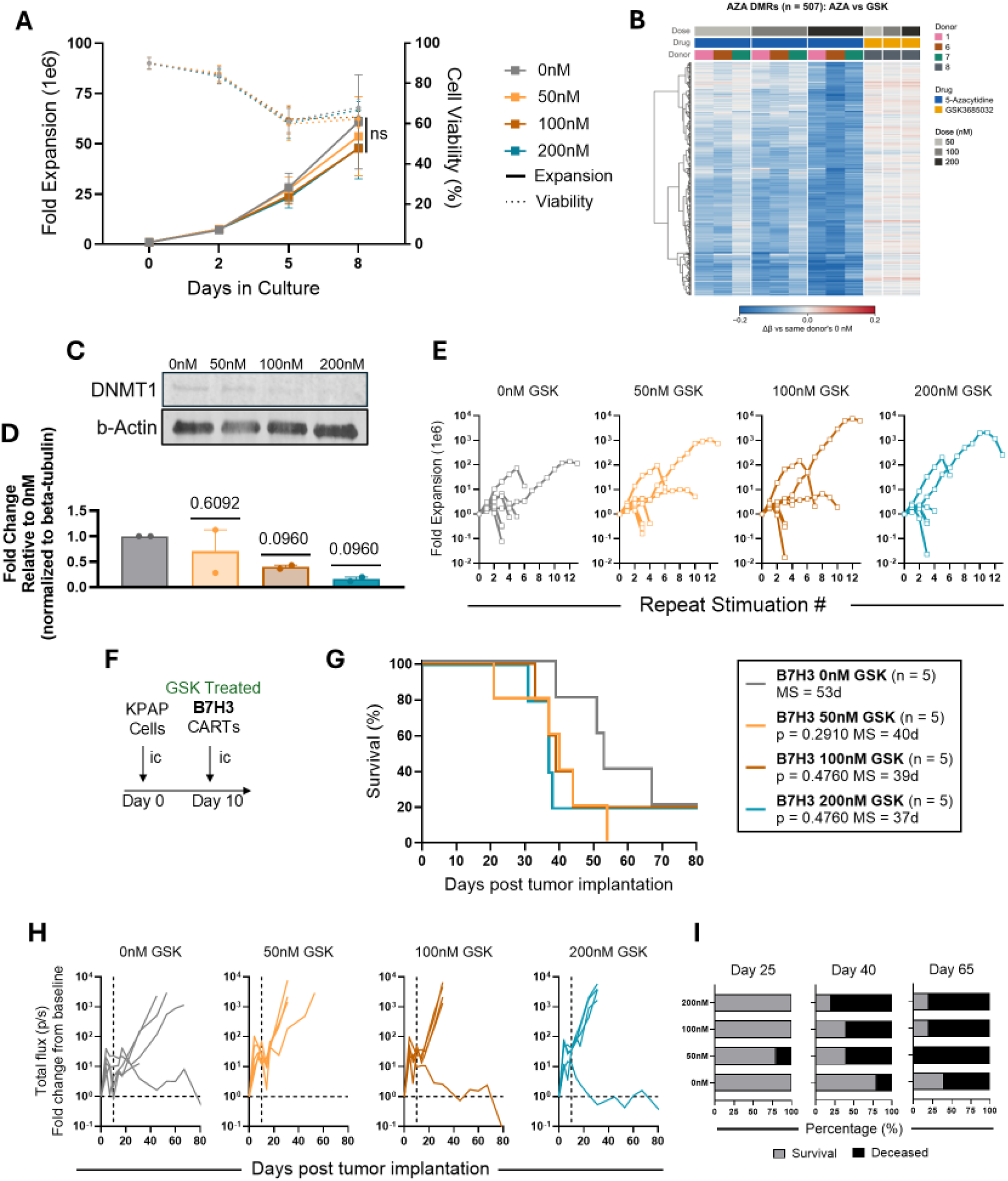
Functional Assessment of GSK365032 Treated CAR T-cells *in vitro* and *in vivo.* **A)** CAR T cells were expanded with IL-2 (50 U/mL) for 8 days following transduction and fold expansion with viability were monitored. (mean ± SEM, n = 12 donors, two-way repeated-measures ANOVA with Dunnett’s multiple-comparisons test comparing each treatment group to the 0 nM control at each time point (n = 6). **B)** Heatmap of methylation changes following AZA and GSK3685032 treatment across 507 AZA-defined DMRs. DMRs were identified by dose-associated hypomethylation following AZA treatment (DMRcate FDR < 0.05; |mean Δβ, 0→200 nM| > 0.10; n = 12 samples, 3 donors × 4 doses). Δβ represents methylation change at 50, 100, and 200 nM relative to the corresponding donor’s 0 nM condition, averaged across probes within each DMR. Rows are clustered using Ward’s method; columns are ordered by treatment and dose and annotated by drug, donor, and dose.**C-D)** Protein was extracted from CAR T-cells 6 days post transduction and immunoblots were run for DNMT1 with α-tubulin as a loading control. **C)** Representative cropped blots are shown here, and black lines indicate where one part of the blot ends and another begins. **D)** DNMT1 expression was normalized to the corresponding untreated (0nM) control. Treatment conditions were compared with the theoretical value of 1 using two- sided one-sample t-tests with Holm correction for all pairwise multiple comparisons, (n = 2, mean ± SEM). **E)** Following 5- Aza treatment, B7H3 CAR T cells were serially challenged with GL261 tumor cells at a 2:1 effector-to-target ratio and restimulated every 2-3 days with fresh targets until proliferative expansion and/or cytotoxic activity could no longer be sustained. Fold expansion across successive stimulation rounds is shown for individual donors (n = 7). **F-J)** 1e5 KPAP.ffLuc cells were injected orthotopically into Albino C57BL/6 mice, followed by intra-tumoral injection of 1e6 B7H3 CAR T-cells. **F)** Experimental scheme. **G)** Kaplan-Meier survival curve (n =5, log-rank Mantel-Cox test with Holm-Šídák correction for multiple comparisons to 0nM, MS = median survival). **H)** Quantification of tumor burden over time as fold change in bioluminescent flux relative to baseline imaging. **I)** Summary plots showing percentage of surviving and deceased mice within each treatment group at days 25, 40 and 65 after tumor implantation.

To assess CAR T-cell phenotype and immune composition following treatment, KPAP tumors were harvested 4 days after CAR T-cell administration and analyzed by high-dimensional flow cytometry (Supplementary Figure 11). A numerically greater frequency of CAR⁺ (CD90.1⁺) cells was detected in tumors from mice treated with 50nM 5-Aza-primed CAR T-cells compared with the 0nM (*p*=0.2013) and 100nM (*p*=0.2014) groups (Figure 4H). Phenotypic analysis of intratumoral CAR⁺ cells revealed increased expression of activation and effector-associated markers, including CD25, IFNγ, and TNFα, in the 50nM group, whereas expression of these markers was reduced in the 100nM group (Figure 4I). CAR T-cells manufactured in the presence of 5-Aza also exhibited a general reduction in regulatory-associated markers, including AHR, IL-10, and PD-1, following *in vivo* administration.

### 5-Aza-primed B7H3 CAR T-cells demonstrate enhanced efficacy in syngeneic ovarian cancer

To determine whether the benefits of 5-Aza priming extended beyond intracranial tumors, we evaluated B7H3 CAR T-cells in the syngeneic ID8 Trp53^−/-^ ovarian cancer model^33^. ID8 cells were implanted into the peritoneal cavity of female mice, followed by intraperitoneal injection of 3e^6^ B7H3 CAR T-cells three weeks later (Figure 4J). The survival benefit of 5-Aza priming was most pronounced at 50nM (Figure 4K), with a median survival of 59 days and prolonged survival relative to 0nM CAR T-cells (*p* = 0.0763), whereas 0nM and 100nM treated CAR T-cells produced similar survival outcomes (median survival, 50 days; *p* > 0.05), similar to observations in the brain tumor models. Mice receiving 50nM primed CAR T-cells exhibited delayed tumor associated weight gain (Figure 4L) and a significant delay in ascites development (tumor burden) after 49 days compared to 0nM and 100nM (*p* = 0.0206), with continued control at day 67 (Figure 4M). Control CAR T-cells did not improve survival, regardless of 5-Aza exposure compared to tumor only control (all *p*>0.05; Supplementary Figure 12A).

### GSK3685032 priming during CAR T-cell manufacturing does not induce functional benefit

To determine whether selective DNMT1 inhibition could reproduce the effects observed with 5-Aza, we next evaluated GSK3685032 (GSK) during CAR T-cell manufacturing. GSK-treated CAR T-cells were manufactured using the same concentrations and treatment timeline employed for 5-Aza-primed CAR T-cells. GSK treatment did not affect B7H3 CAR T-cell viability or expansion following manufacturing (Figure 5A).

To evaluate the epigenetic remodeling of selective DNMT1 inhibition, GSK-treated B7H3 CAR T-cells were profiled using the Infinium Mouse Methylation BeadChip. GSK-treated CAR T-cells showed comparatively limited methylation changes across the 507 Aza-defined DMRs that exhibited dose- associated hypomethylation (Figure 5B). Additionally, GSK-treated samples showed fewer nominally significant differentially methylated probes (∼15,00 vs 47,000-86,000). Consistent with a limited genome-wide methylation effect, median Δβ remained near zero across GSK concentrations, whereas Aza-treated donors exhibited progressively greater hypomethylation with increasing dose (Supplementary Figure 13). Despite this, GSK induced a dose-dependent reduction in DNMT1 protein abundance (Figure 5C-D), with DNMT1 levels reduced to 0.70-, 0.40-, and 0.16-fold relative to untreated controls in the 50, 100, and 200nM groups, respectively. GSK treatment had minimal effects on CAR T-cell transduction or phenotype, with no significant differences in CD8 frequency, memory subset distribution, or inhibitory receptor expression (Supplementary Figure 14A-D).

During serial GL261 coculture, GSK did not significantly alter maximum expansion or cumulative tumor killing (Figure 5E; Supplementary Figure 14E-F). MTS assays showed no significant differences in cytotoxicity, with early separation during the third stimulation becoming variable and inconsistent by the sixth stimulation (Supplementary Figure 11G). At the fourth stimulation, IL-2 secretion was reduced in GSK-treated groups, whereas IFNγ secretion increased and reached significance at 100nM (*p*=0.0192), a cytokine profile distinct from that observed with 5-Aza (Supplementary Figure 14H).

Finally, we evaluated the antitumor activity of GSK-primed B7H3 CAR T-cells in the orthotopic, immunocompetent KPAP DMG model. Mice received 1e⁶ CAR T-cells intratumorally on day 10 after implantation (Figure 5F). Untreated B7H3 CAR T-cells demonstrated the most durable tumor control, with delayed increases in bioluminescent signal and preservation of body weight relative to GSK- treated groups (Figure 5G-I; Supplementary Figure 15). Non-treated CAR T-cells also provided the greatest improvement in median survival with 53 days compared with 29 days for tumor-only controls and 40, 39, and 37 days for the 50, 100, and 200nM GSK groups, respectively. All conditions significantly improved survival relative to tumor-only controls (p<0.05, Supplemental Figure 10A), but none improved survival relative to untreated CAR T-cells.

### Therapeutic benefits of 5-Aza priming extend to an alternative CAR T-cell construct

Having demonstrated reproducible improvements in B7H3 CAR T-cell efficacy across multiple tumor models and finding that selective DNMT1 inhibition did not recapitulate these effects, we next investigated whether the benefits of 5-Aza extended to an alternative CAR platform. GD2-targeted CAR T-cells were manufactured in the presence of 5-Aza using the same production strategy described above.

Comparable transduction efficiencies were achieved across treatment groups, with a mean GD2 CAR transduction efficiency of 68% (SD ±34.2%), and phenotypic characterization identified no major differences prior to administration (Supplementary Figure 16A-D). Annexin V-based assays using KPAP cells confirmed GD2-specific cytotoxicity and identified modest improvements in tumor elimination among 5-Aza-primed CAR T-cell products *in vitro* (Supplementary Figure 16E-G).

We next evaluated the antitumor efficacy of 5-Aza-treated GD2 CAR T-cells *in vivo* with the KPAP model (Figure 6A). The 100nM group showed the greatest tumor control, with reduced bioluminescent signal and a median survival of 95 days, compared with 39 and 37 days for the 0nM and 50nM groups, respectively (Figure 6B–C; Supplementary Figure 17). Despite the longer survival observed with 100nM treatment, survival did not significantly differ from non-treated CAR T-cells (p=0.2650; Figure 6B). All GD2 CAR T-cell groups significantly prolonged survival relative to tumor- only controls (Supplementary Figure 10B).

**Figure 6.**
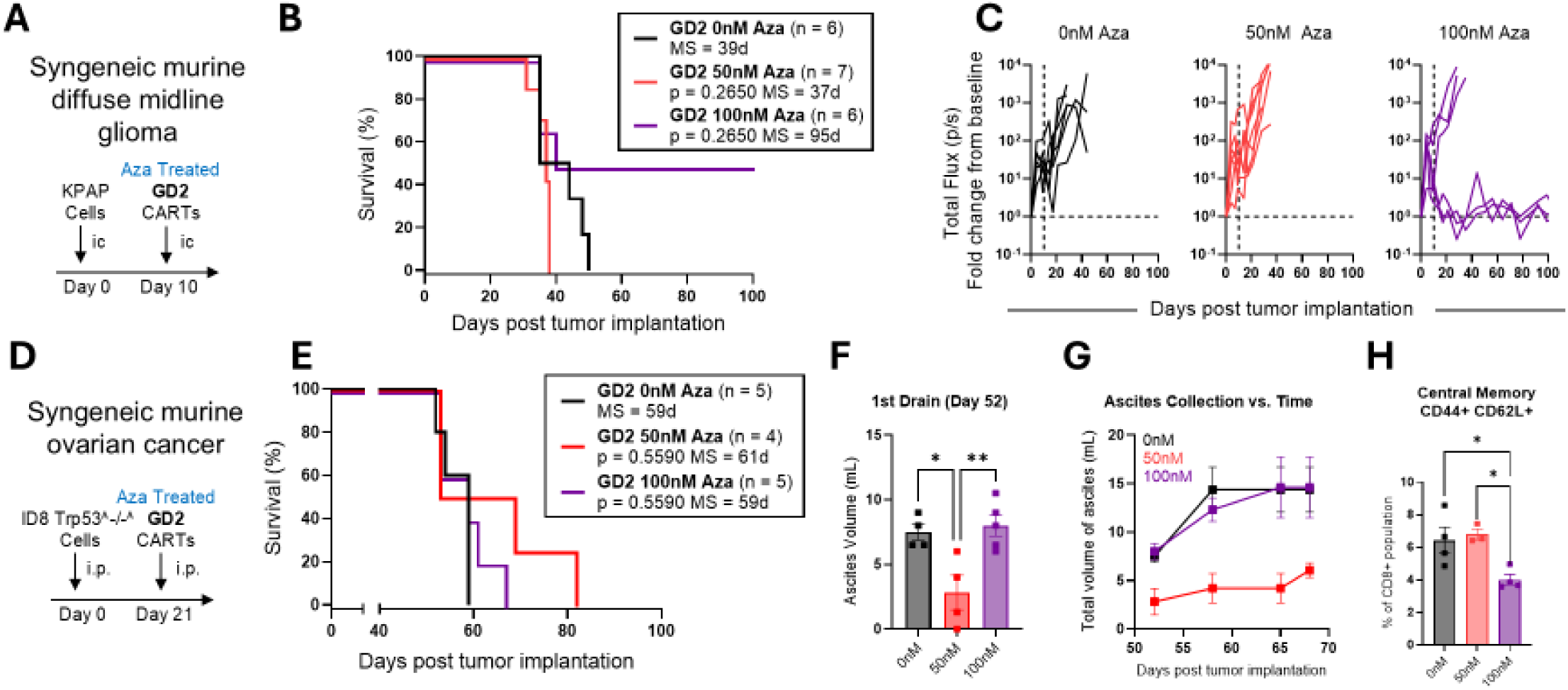
Evaluation of 5-Aza treated GD2 CAR T cells *in vivo*. **A-C**) 1e5 KPAP.ffLuc cells were injected orthotopically into Albino C57BL/6 mice, followed by intra-tumoral injection of 2e6 B7H3 CAR T-cells. **A)** Experimental scheme. **B)** Kaplan-Meier survival curve (n = 6-7, log-rank Mantel-Cox test with Holm-Šídák for multiple comparisons to 0nM, MS = median survival). **C)** Quantification of tumor burden over time as fold change in bioluminescent flux relative to baseline imaging. **D-L)** 5e6 ID8 *Trp53-/-* murine OC were injected intraperitoneally into C57BL/6 mice, followed by intraperitoneal injection of B7H3 CAR T cells. **D)** Experimental scheme. **E)** Kaplan-Meier survival curve (n = 4-5, log-rank Mantel-Cox test with Holm-Šídák correction for multiple comparisons to 0nM). One mouse in 50nM treatment group was excluded from analysis due to failure to demonstrate tumor establishment, as no increase in abdominal circumference or subsequent tumor-associated mortality was observed. **F)** Volume of ascites collected from treated mice at day 52; 1^st^ drain (n = 5, repeated-measures one-way ANOVA with Tukey’s multiple-comparisons test, *P < 0.05, **P < 0.01). **G)** Cumulative total volume of ascites collected over time. **H)** Frequency of effector memory (CD44+CD62L+) CD90.1+ CAR T cells recovered from ascites (n = 3-4, repeated-measures one-way ANOVA with Tukey’s multiple-comparisons test, *P < 0.05).

### 5-Aza priming delays ovarian cancer progression across B7H3 and GD2 CAR T-cell platforms

To determine whether the benefits of 5-Aza priming extended across CAR constructs and tumor models, we next evaluated GD2 CAR T-cells in the syngeneic ID8-Trp53^−/−^ ovarian cancer model (Figure 6D). Survival differences among CAR T-cell treatment groups did not reach statistical significance, although mice treated with 50nM 5-Aza-primed GD2 CAR T-cells achieved improved median survival (61 days vs 59 days for the 0 and 100nM groups; Figure 6E). The 50nM group exhibited significantly reduced ascites burden at first drains compared with both the 0nM (*p*=0.0195) and 100nM (*p*=0.0080) groups and delayed ascites accumulation throughout the study (Figure 6F-G; Supplementary Figure 16A-B). Compared with tumor-only controls, non-treated CAR T-cells provided minimal survival benefit, whereas both 50nM and 100nM 5-Aza-treated CAR T-cells extended survival, with the 100nM group reaching statistical significance (*p*=0.0473; Supplementary Figure 12B).

To further characterize the tumor-associated immune compartment, cells recovered from the first ascites drains were analyzed by flow cytometry. Mice treated with 100nM-primed CAR T-cells exhibited reduced frequencies of CD44⁺CD62L^+^ central memory CD8⁺ T-cells relative to the 50nM (*p*=0.0255) and 0nM (*p*=0.0343) groups (Figure 6H), with additional modest differences in immune populations (Supplementary Figure 16C-D).

## DISCUSSION

Epigenetic regulation contributes to T-cell differentiation and functional state^34,35^, providing a rationale for targeting DNA methylation during CAR T-cell manufacturing. Genetic disruption of epigenetic regulators has demonstrated the potential to improve T-cell expansion, persistence and function^21,36,37^. However, incorporating additional genetic modifications into CAR T-cell engineering workflows presents challenges for clinical translation^38^. We therefore investigated transient low-dose pharmacologic DNMT inhibition during CAR T-cell manufacturing as an alternative strategy to enhance CAR T-cell function against solid tumors. Murine CAR T-cells were manufactured in the presence of 5-azacytidine (5-Aza) or GSK3685032 (GSK). Transient low-dose 5-Aza exposure produced durable functional benefits, including enhanced expansion during prolonged serial antigen challenge and improved antitumor activity across multiple immunocompetent syngeneic tumor models and CAR constructs. In contrast, selective DNMT1 inhibition with GSK did not recapitulate these benefits, suggesting that the functional effects of 5-Aza may be driven predominantly by DNMT3A inhibition^21^ rather than DNMT1 inhibition alone. These findings demonstrate that transient pharmacologic modulation of DNA methylation during CAR T-cell manufacturing can produce durable functional changes that persist beyond the treatment window.

5-Aza-treated CAR T-cells demonstrated dose-associated epigenetic and transcriptional remodeling after manufacturing. Although direct overlap between differentially methylated and expressed genes was limited, pathway-level analyses identified concordant changes in lymphocyte-associated programs, suggesting broader remodeling rather than specific remodeling of a limited set of genes. Dose-associated expression changes in T-cell effector genes further supported this pattern. Interestingly, these molecular changes occurred without substantial alterations in the overall phenotypic composition of the manufactured CAR T-cell product, suggesting that 5-Aza did not broadly redirect T- cell differentiation during manufacturing. Instead, transient epigenetic remodeling may alter the capacity of CAR T-cells to respond to subsequent antigenic stimulation. Consistent with this interpretation, IL-2 secretion was initially reduced following the first tumor challenge but increased relative to untreated CAR T-cells with subsequent stimulation, suggesting an altered response to repeated antigen exposure rather than immediate enhancement of effector function. Given the role of sustained IL-2 signaling in supporting T-cell function^39,40^, this response may contribute to the sustained functional capacity of 5- Aza-treated CAR T-cells.

Despite dose-dependent molecular remodeling following 5-Aza exposure, increasing drug concentration did not uniformly translate to greater antitumor activity, suggesting that effective epigenetic priming may occur within an optimal dose range. This was particularly evident in immunocompetent *in vivo* models, where 50nM 5-Aza consistently enhanced B7H3 CAR T-cell activity across both brain and ovarian cancer models, improving survival and delaying tumor progression. In the KPAP model, this benefit was accompanied by greater CAR T-cell abundance within the brain after treatment, suggesting increased persistence within the tumor-bearing tissue. In contrast, despite promising functional activity at 100nM, B7H3 CAR T-cells manufactured at this dose were associated with early mortality in the KPAP model without a corresponding increase in bioluminescent tumor burden. Given the potent antitumor activity of the B7H3 CAR, more extensive epigenetic modulation may contribute to treatment-associated toxicity, although the underlying mechanism remains unclear. Similar treatment-associated toxicities following robust CAR T-cell activation have been reported previously, providing biological precedent for this possibility^41^.

Notably, this dose relationship differed by both CAR construct and tumor model. In KPAP, 100nM 5- Aza-treated GD2 CAR T-cells improved survival relative to the 50nM condition, contrasting with the response of B7H3 CAR T-cells. Conversely, in the aggressive ovarian tumor model, 50nM 5-Aza enhanced therapeutic activity with both CAR constructs. Although the magnitude of improvement appears modest, this finding is notable in the context of the aggressive ID8 model, where B7H3 and GD2 CAR T-cell treatment alone provided limited survival benefit over tumor-only controls. Thus, even modest enhancement of antitumor activity following 5-Aza priming may be meaningful in this therapeutically challenging setting. These differences suggest that optimal DNMTi exposure depends on both CAR construct and disease context, supporting product-specific rather than uniform dosing.

In contrast to 5-Aza, GSK-mediated DNMT1 depletion was not accompanied by dose-dependent changes in DNA methylation or functional benefit. GSK-treated CAR T-cells demonstrated limited activity across *in vitro* assays and, notably, reduced survival relative to untreated CAR T-cells *in vivo*. These findings suggest that suppression of DNMT1 alone is insufficient to reproduce the beneficial effects of 5-Aza and may negatively affect CAR T-cell function. Whereas 5-Aza incorporates into replicating DNA and traps engaged DNMTs, GSK selectively inhibits DNMT1, resulting in more direct and widespread degradation of the cellular DNMT1 protein^42^. This distinction may be particularly relevant given the emerging noncanonical functions of DNMT1, including regulation of gene expression independently of its catalytic activity and participation in cell-cycle regulation, DNA damage repair, and stem cell function^19,43^. These findings demonstrate that pharmacodynamic inhibition of DNMT1 is not itself predictive of improved CAR T-cell function and underscore the importance of the mechanism by which epigenetic modulation is achieved during manufacturing.

The use of murine CAR T-cells enabled evaluation in syngeneic, immunocompetent tumor models; however, a limitation of this study is that the optimal 5-Aza exposure identified here may not directly translate to human CAR T-cell manufacturing. Future studies will therefore require optimization of 5- Aza dosing for human CAR T-cells and evaluation of whether these functional and epigenetic effects are conserved across species, CAR constructs, and disease settings. Our findings support pharmacologic epigenetic modulation during *ex vivo* manufacturing as a clinically adaptable strategy to enhance CAR T-cell function across brain and ovarian cancer models. More broadly, this study supports transient small-molecule modulation as a potentially versatile approach for improving CAR T-cell products during manufacturing.

## Acknowledgements

The authors are grateful to the Integrated Biomedical Sciences program at The George Washington University for its continued support of graduate student training and to the Research Animal Facility at Children’s National Medical Center and George Washington University for their assistance and support throughout the animal studies. We sincerely thank the Brain Tumor Institute Bioinformatics Core at the Children’s National Research Institute (CNRI) for their bioinformatics guidance and the Flow Cytometry Core for their continued collaboration through the Cell Enhancement and Technologies for Immunotherapy Program (CETI) at Children’s National. The authors are grateful to Dr Iain McNeish for providing the 123 isogenic Trp53−/− murine ovarian cancer cell line, Dr. Zoe Weaver Ohler for providing the MBR525-1 murine glioblastoma cell line and Dr Cheng-Chia “Fred” Wu for providing the KPAP murine diffuse midline glioma cell line. We also thank the Genome Technology Access Center at the McDonnell Genome Institute at Washington University for their assistance with sequencing. Finally, the authors extend their gratitude to William Bishop for his contributions to this work as a student intern.

