## Supplemental Figures for "DNA Methyltransferase Inhibition During CAR T-cell Manufacturing Enhances Antitumor Activity in Immunocompetent Brain and Ovarian Cancer Models"

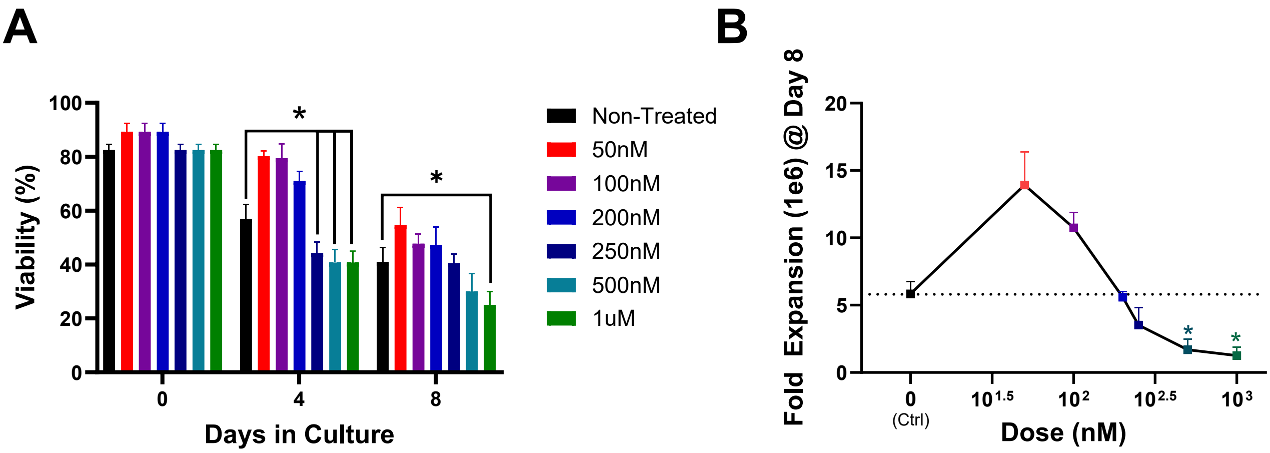


**Supplemental Figure 1. 5-Aza dose titration to evaluate CAR T cell viability and expansion.** CAR T cells were expanded with IL-2 (50 U/mL) for 8 days following transduction. 5-Aza treatment was suspended after 3 days of treatment and fold expansion with viability were monitored throughout culture. Data are shown as mean ± SEM (n = 4 donors). **A)** Viability over time was analyzed by two-way repeated-measures ANOVA with Dunnett's multiple-comparisons test comparing each treatment group to the non-treated control (0nM) at each time point, *P < 0.05. group, analyzed by one-way ANOVA with Dunnett's multiple-comparisons test comparing expansion of each treatment group to the non-treated control, *P < 0.05.

.

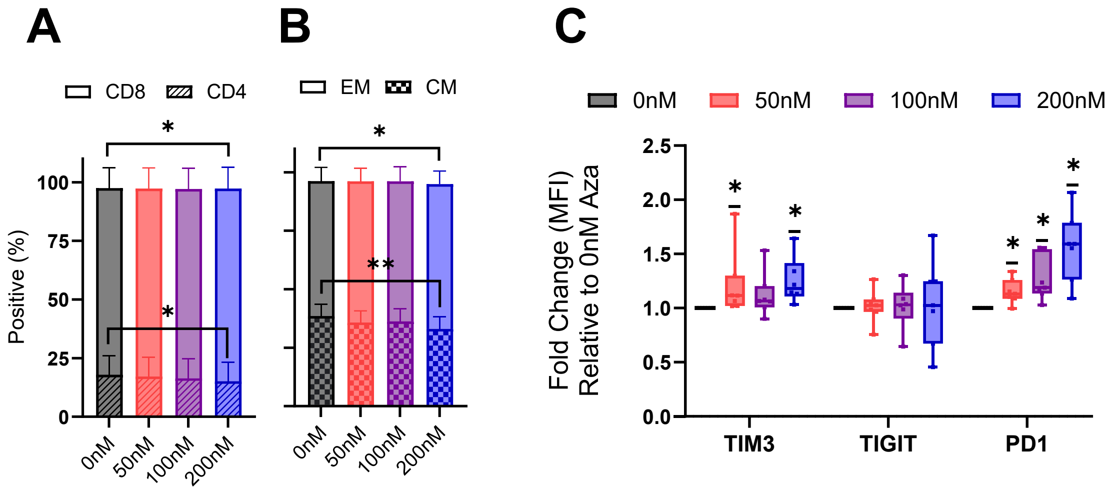


**Supplemental Figure 2. 5-Aza treated B7H3 CAR T-cell phenotype following manufacturing.** At day 6 post-transduction, CAR T-cells were analyzed by flow cytometry for phenotypic and exhaustion markers. **A)** CD4 and CD8 composition and **B)** effector memory (EM; CD44+/CD62L−) and central memory (CM; CD44+/CD62L+) phenotype distribution of the CD8+ population. Data are shown as mean ± SEM (n = 10) and were analyzed by two-way ANOVA with Tukey's multiple-comparisons test comparing treatment conditions within each phenotype, *P < 0.05, **P < 0.01. **C)** Fold change in exhaustion marker median fluorescence intensity (MFI) at each 5-Aza concentration, normalized to the donor-matched non-treated control (0 nM). Data are shown as minimum-to-maximum range (n = 7) and were analyzed by one-sample Wilcoxon signed-rank test against a hypothetical value of 1.0, *P < 0.05.

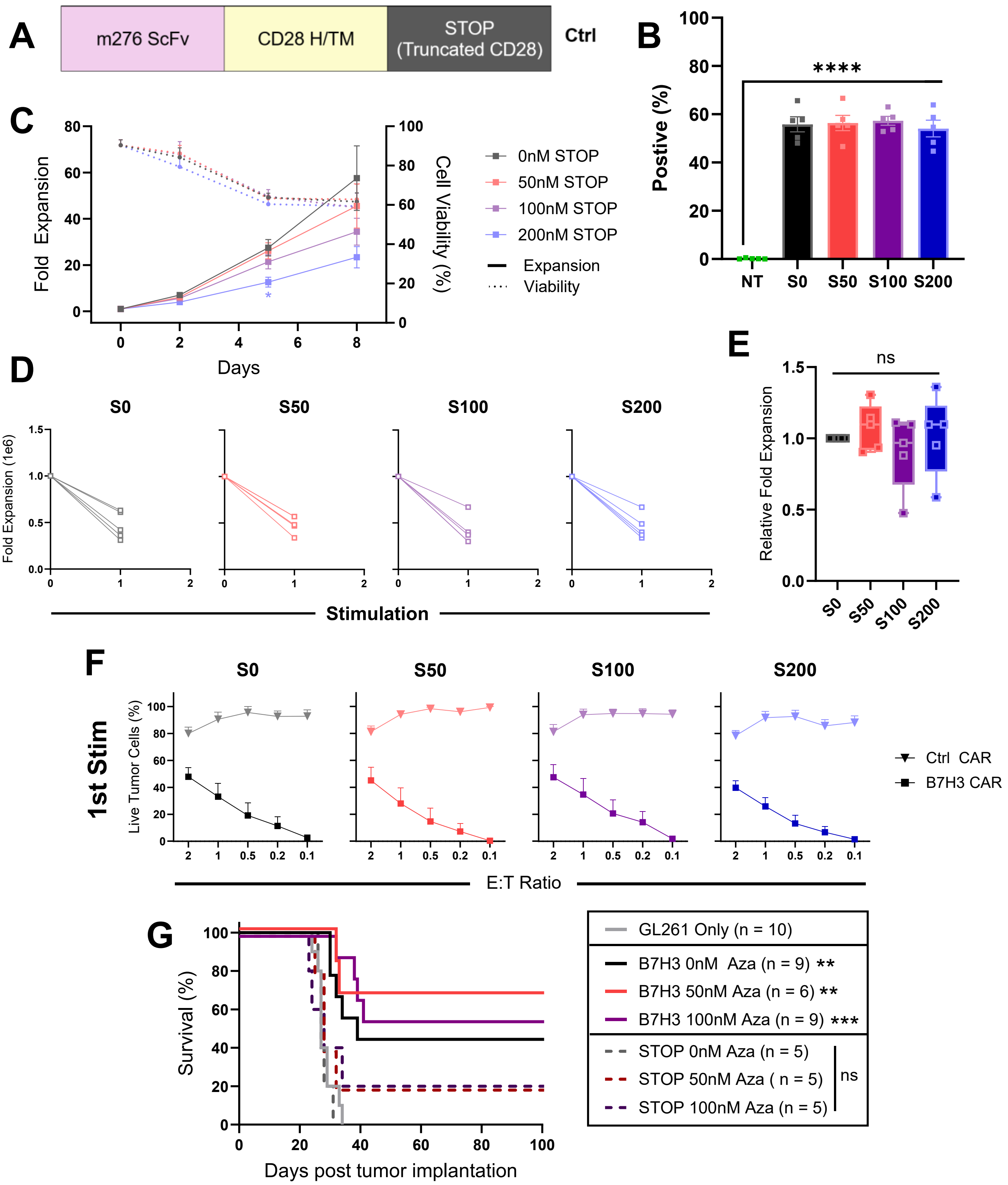


**Supplemental Figure 3. Characterization of 5-Aza-treated CD28 truncated Ctrl CAR T cells. A)** Schematic of the Ctrl CAR construct. **B)** CAR expression evaluated by flow cytometry on days 5–6 post-transduction. Representative flow plots of non-transduced (NT) and Ctrl CAR murine T cells and summary of F(ab')$_2$-positive T cells (n = 5, mean ± SEM, one-way ANOVA with Tukey's multiple-comparisons test, ****P < 0.0001). **C)** Ctrl CAR T cells were expanded with IL-2 (50 U/mL) for 8 days following transduction. Fold expansion and viability were monitored throughout culture and are shown as mean ± SEM (n = 5 donors). Expansion kinetics were analyzed by two-way repeated-measures ANOVA with Dunnett's multiple-comparisons test comparing each treatment group with the 0 nM control at each time point, , *P < 0.05. **D)** Following 5-Aza treatment, Ctrl CAR T cells were challenged with GL261 tumor cells. Ctrl CAR T cells did not achieve sufficient tumor cell killing to permit subsequent restimulation; therefore, fold expansion following a single stimulation is shown for individual donors (n = 5). **E)** Fold expansion following a single stimulation at each 5-Aza concentration relative to the donor-matched untreated control, calculated as the treated maximum divided by the 0 nM maximum (n = 5, minimum-to-maximum range, one-sample Wilcoxon signed-rank test versus a hypothetical value of 1.0). **F)** MTS cytotoxicity assay of Ctrl (triangle) and B7-H3 (square) CAR T cells against GL261 tumor cells across indicated effector-to-target (E:T) ratios following the first and only stimulation (n = 5, mean ± SEM). **G)** C57BL/6 mice received orthotopic injection of 1 × 10⁵ GL261 cells followed by intratumoral administration of 5-Aza-treated Ctrl CAR or B7H3 CAR T cells. Kaplan-Meier survival curves are shown (n = 6–10; log-rank Mantel-Cox test with Holm-Šídák correction for multiple comparisons versus GL261 only). Asterisks denote comparisons with the tumor-only control: *P < 0.05, **P < 0.01, ***$P$ < 0.001.

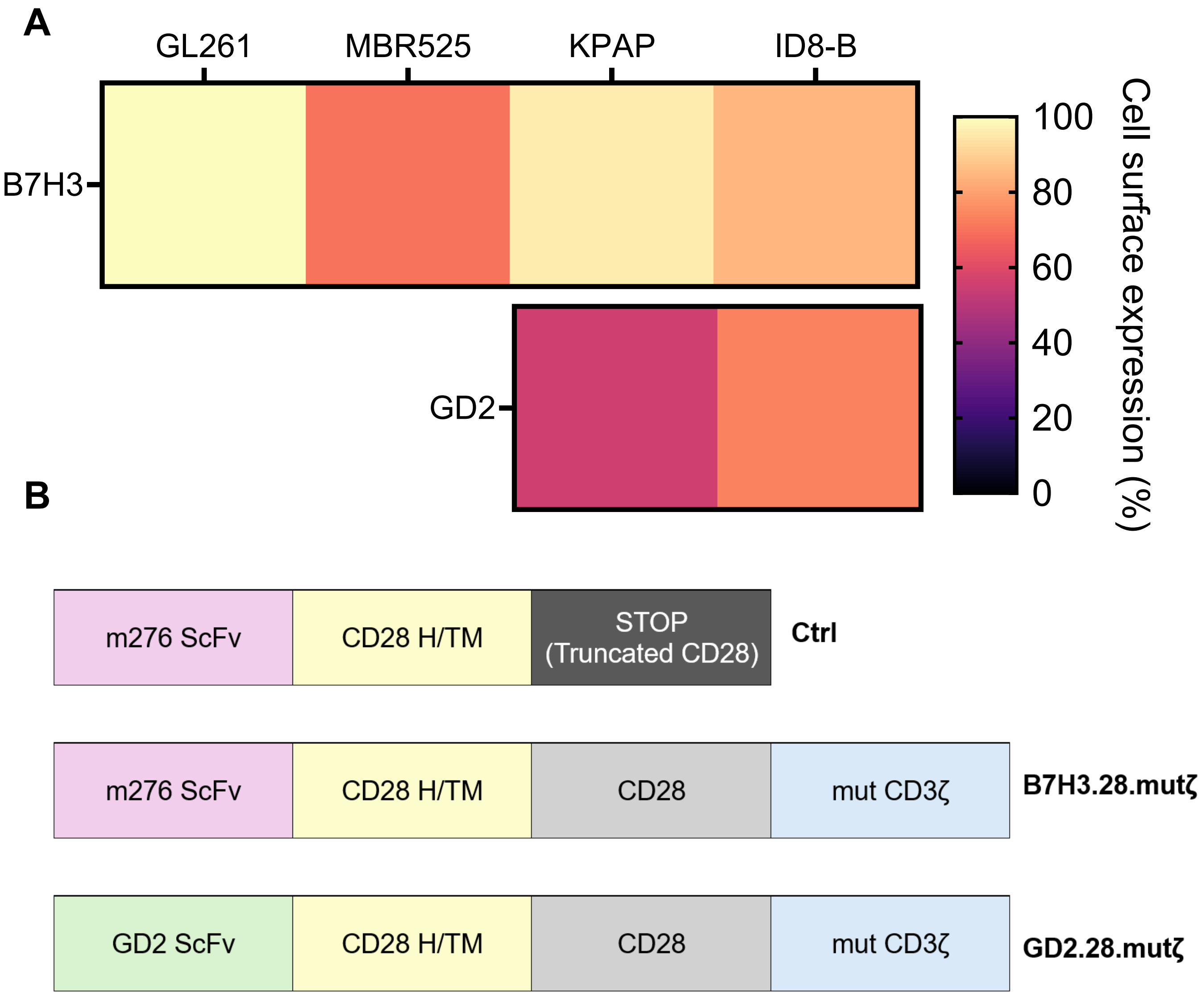


**Supplemental Figure 4. Target antigen expression on utilized cell lines and corresponding CAR constructs. A)** Heatmap showing B7H3 and GD2 cell-surface expression on GL261, MBR525, KPAP, and ID8-B tumor cells, as assessed by flow cytometry. Color represents the percentage of tumor cells positive for each indicated antigen (0–100%). **B)** Scheme of utilized B7H3 and GD2 murine CAR constructs.

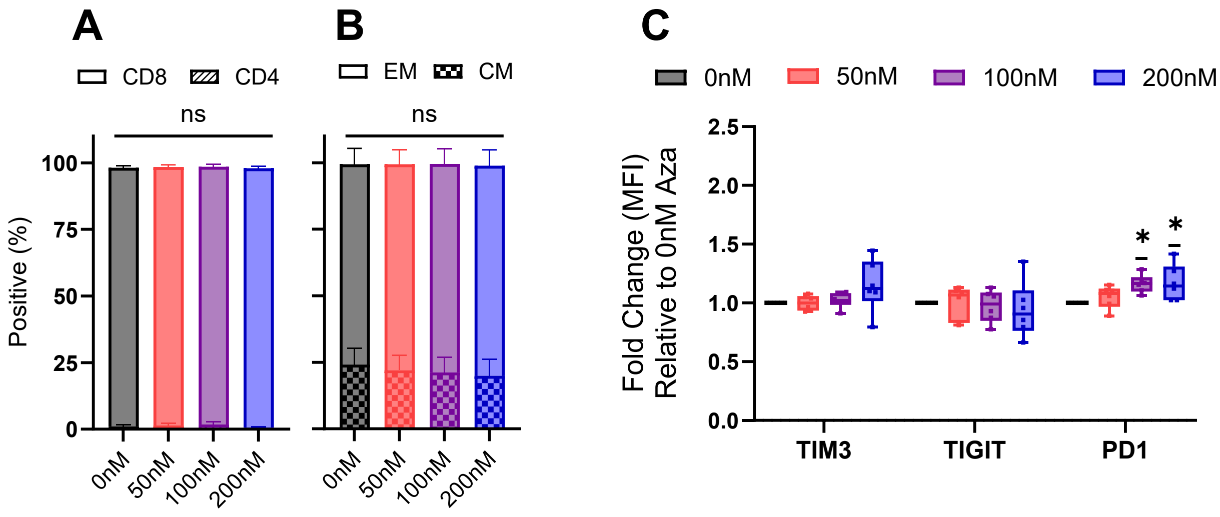


**Supplemental Figure 5. 5-Aza-treated B7H3 CAR T-cell phenotype following three repeat stimulations with B7H3+ GL261.** Following three rounds of repeated stimulation resulting in T-cell expansion and tumor cell elimination, CAR T cells were collected from co-culture and analyzed by flow cytometry for phenotypic and exhaustion markers. **A)** CD4 and CD8 composition and **B)** effector memory (EM; CD44+/CD62L−) and central memory (CM; CD44+/CD62L+) phenotype distribution of the CD8+ population. Data are shown as mean ± SEM (n = 6) and were analyzed by two-way ANOVA with Tukey's multiple-comparisons test comparing treatment conditions within each phenotype. **C)** Fold change in exhaustion marker median fluorescence intensity (MFI) at each 5-Aza concentration, normalized to the donor-matched non-treated control (0 nM). Data are shown as minimum-to-maximum range (n = 6) and were analyzed by one-sample Wilcoxon signed-rank test against a hypothetical value of 1.0, *P < 0.05.

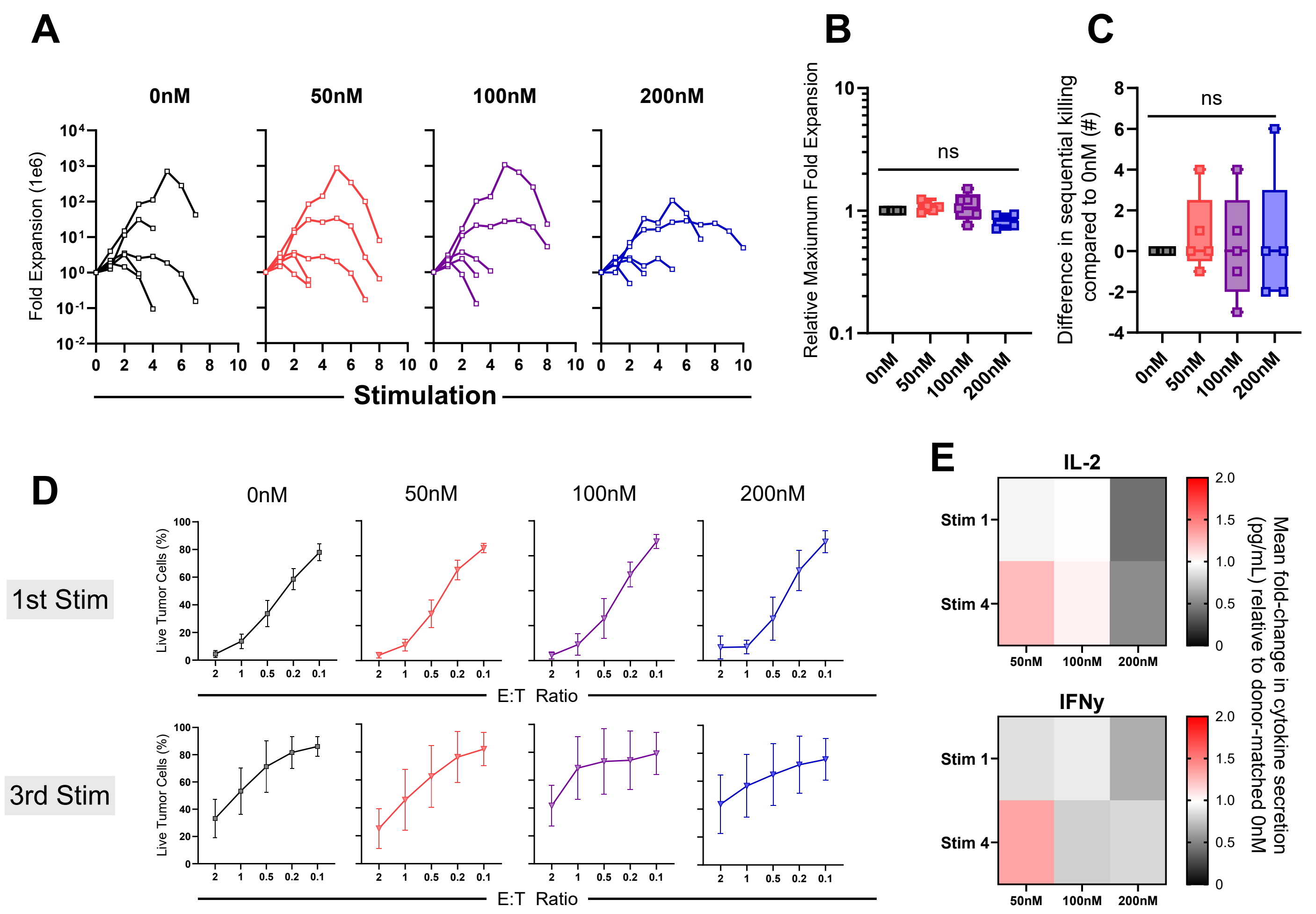


**Supplemental Figure 6. Functional characterization of 5-Aza-treated B7H3 CAR T cells against the MBR525 HGG model. A)** Following 5-Aza treatment, B7H3 CAR T-cells were serially challenged with adherent MBR525 tumor cells at a 2:1 effector-to-target (E:T) ratio and restimulated every 2–3 days with fresh targets until proliferative expansion and/or cytotoxic activity could no longer be sustained. Fold expansion across successive stimulation rounds is shown for individual donors (n = 5). **B)** Maximum fold expansion achived during co-culture at each 5-Aza concentration relative to the donor-matched untreated (0 nM) control, calculated as the treated maximum divided by the 0nM maximum (n = 5, minimum-to-maximum range, one-sample Wilcoxon signed-rank test versus a hypothetical value of 1.0). **C)** Difference in the maximum number of sequential MBR525 tumor-cell stimulations supported while maintaining CAR T-cell cytotoxic activity relative to the donor-matched untreated (0nM) control (n = 5, minimum-to-maximum range, one-sample Wilcoxon signed-rank test versus a hypothetical value of 0). **D)** MTS cytotoxicity assay against MBR525 tumor cells across indicated E:T ratios following the first and third stimulation rounds. Data are shown as mean ± SEM (n = 4). **E)** IL-2 and IFNγ secretion measured by ELISA over consecutive stimulation rounds. Heatmap values represent the mean fold change in cytokine secretion relative to the donor-matched untreated (0nM) control (n = 2–3).

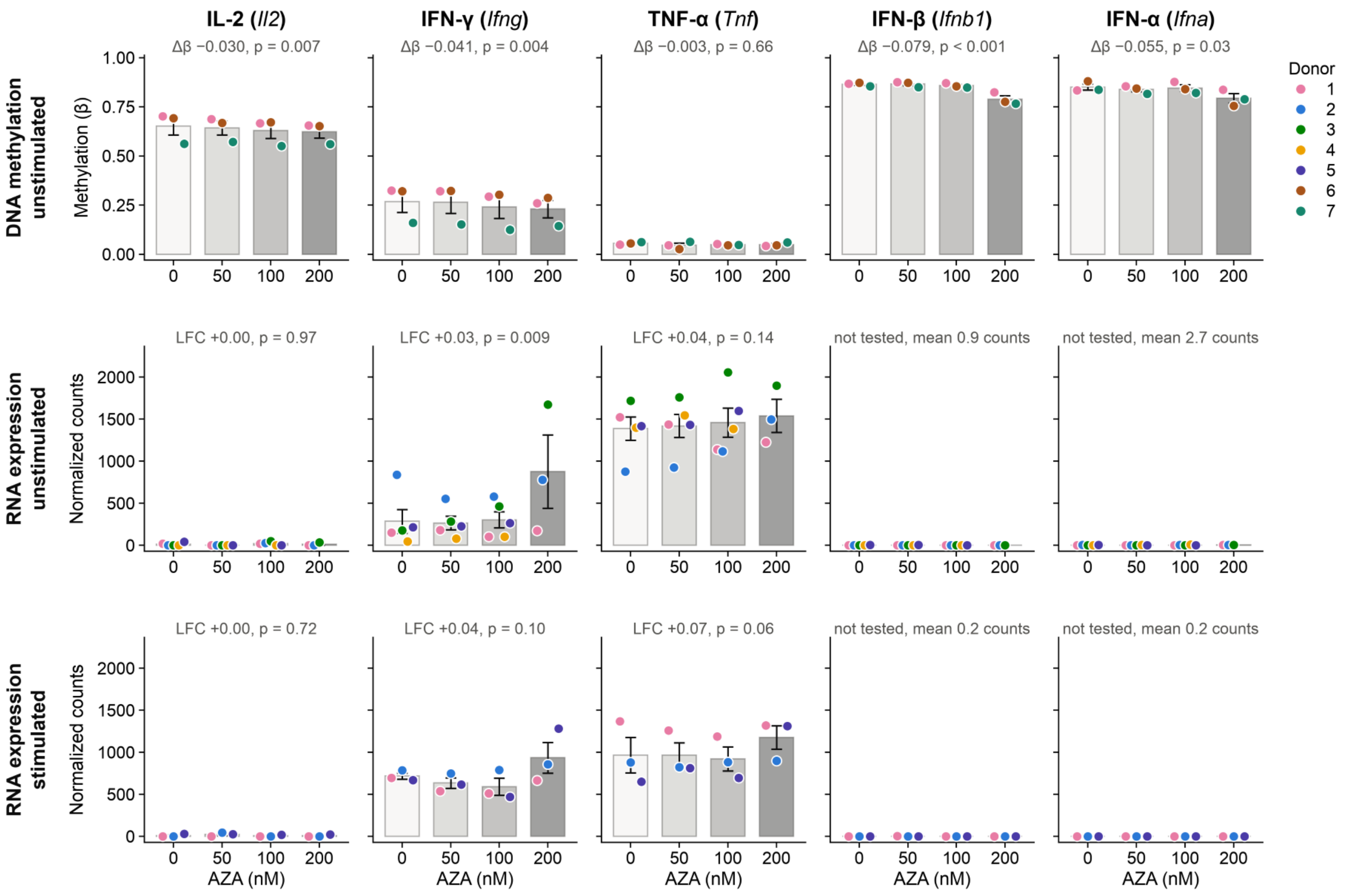


**Supplemental Figure 7. 5-Azacitidine slightly demethylates cytokine loci without significantly changing their expression.** Bars show the mean across donors ± SEM at each dose; points are individual donors. Error bars include baseline differences between donors; dose effects are estimated from donor-blocked models. DNA methylation and RNA expression of IL-2 (*Il2),* IFN-γ (*Ifng*), and TNF-α (*Tnf*) in B7H3 CAR-T cells treated with 5-azacytidine (AZA; 0, 50, 100, 200 nM). Each point is one donor at one dose, shown as absolute β (top row) or normalized counts (middle and bottom rows), and joined across doses. **Top row:** DNA methylation of unstimulated cells (MM285 array; n = 3 donors). Gene value = median β over the gene's promoter (1.5 kb upstream of the TSS) and gene body probes. **Middle row:** RNA expression of unstimulated cells (n = 5 donors). **Bottom row:** RNA expression after stimulation by a single co-culture with GL261-B7H5 tumor cells (n = 3 donors); size factors were estimated jointly across all 30 RNA-seq samples. Labels give the dose effect from donor-blocked linear models with dose as a continuous covariate, fit separately for each condition. p-values are nominal. No RNA effect is significant after genome-wide FDR correction. The methylation decreases at *Il2* and *Ifng* pass a genome-wide FDR test (FDR 0.017 and 0.012), but are no larger than the global AZA-induced loss (median gene Δβ −0.025; 95% of genes lose methylation), and no probes overlap a DMR.

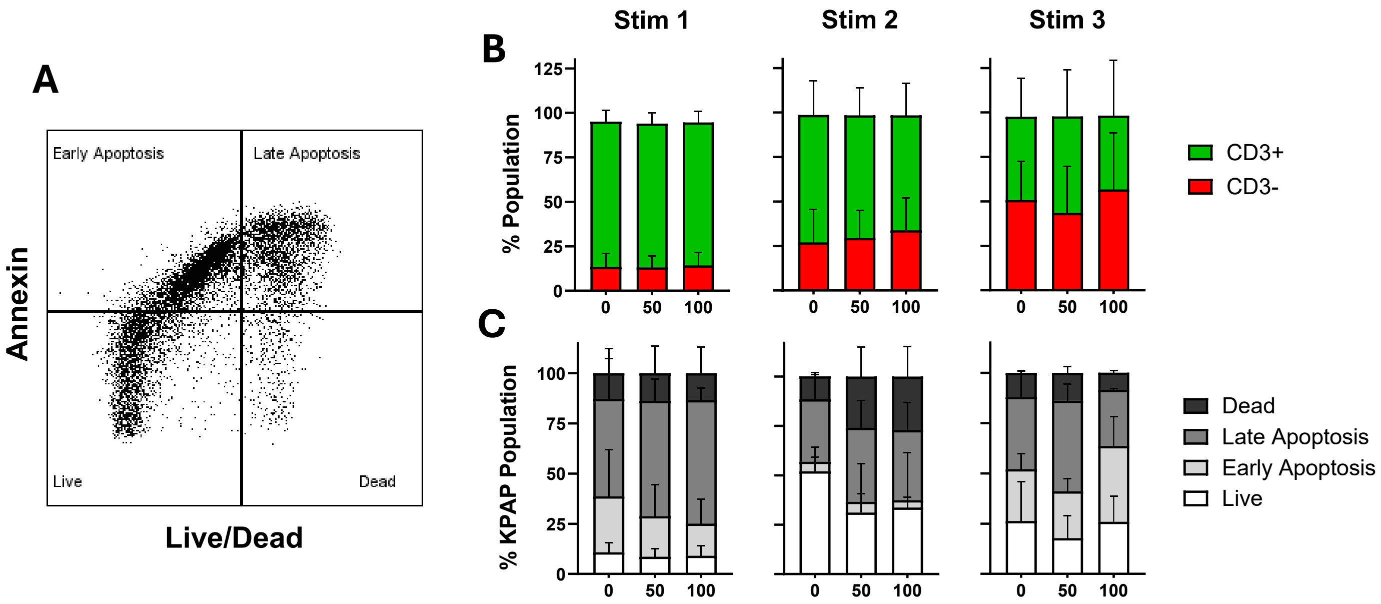


**Supplemental Figure 8. Annexin V-based assessment of B7H3 CAR T-cell cytotoxicity against KPAP. A)** Representative flow cytometric plot of annexin assays conducted on CAR-T cells against suspended KPAP. **B)** The entire collected co-culture was analyzed by flow cytometry to quantify the relative frequencies of CD3+ (CAR-T) cells and CD3− (KPAP) tumor cells. Stacked bar plots show the percentage of each population following co-culture with B7-H3 CAR T cells primed with 0–100 nM 5-Aza across consecutive stimulation rounds (stimulation 1, n = 3; stimulations 2–3, n = 2). **C)** CD3- Tumor cells were classified as live, early apoptotic, late apoptotic, or dead based on Annexin V and live/dead staining. Stacked bar plots show the distribution of tumor cell states following co-culture with B7-H3 CAR T cells primed with 0–100 nM 5-Aza across consecutive stimulation rounds (stimulation 1, n = 3; stimulations 2–3, n = 2).

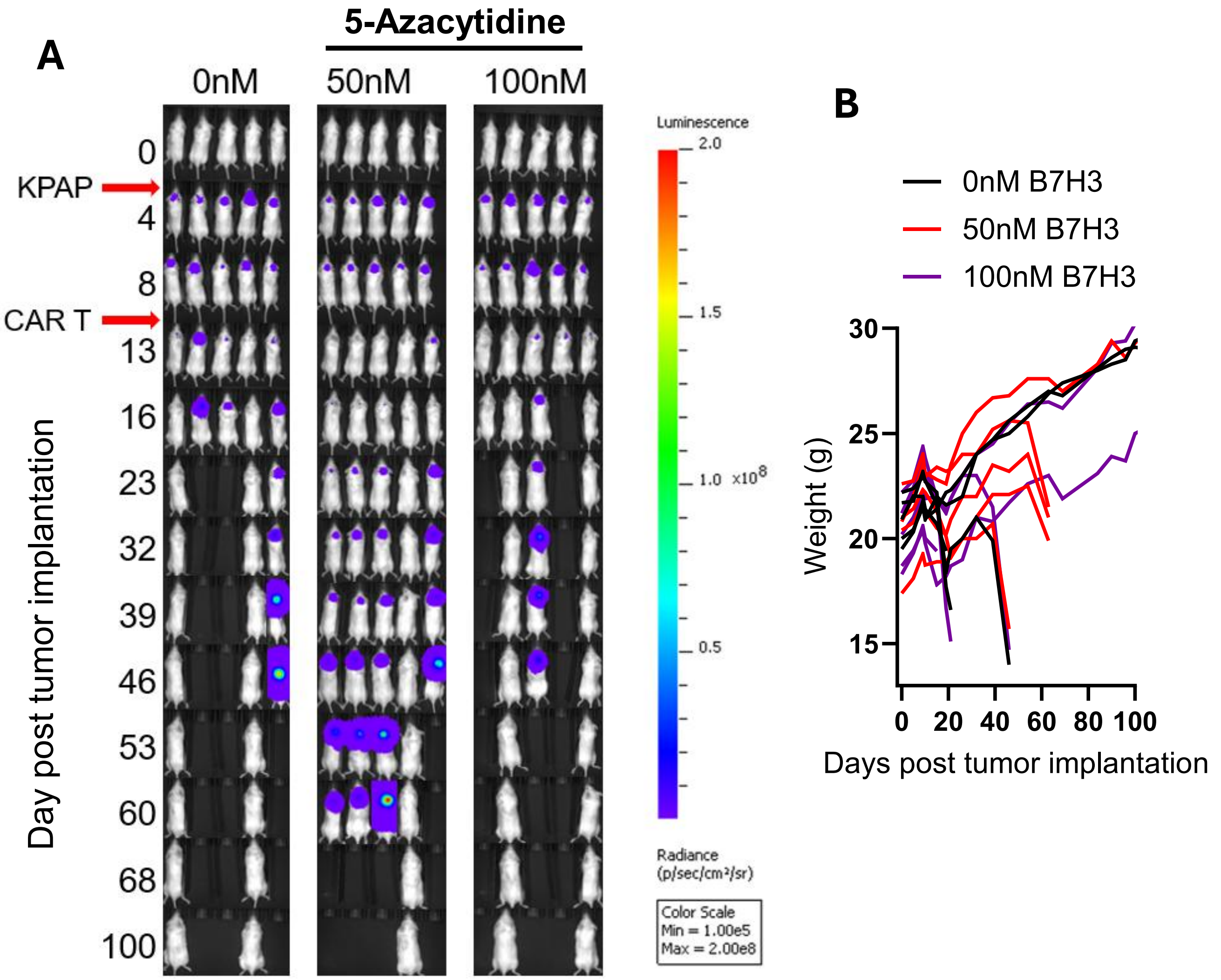


**Supplemental Figure 9. Longitudinal bioluminescent imaging and weight monitoring following 5-Aza-primed B7H3 CAR T-cell therapy in the KPAP model.** On day 0, mice were stereotactically implanted with 1 × $10^5$ KPAP tumor cells. On day 10, mice were treated with 2 × $10^6$ B7-H3 CAR T cells primed with 5-Aza during manufacturing. **A)** BLI images showing tumor burden at the indicated time points following KPAP implantation and B7H3 CAR T-cell treatment. Imaging parameters, shown at left, were kept consistent throughout the study. Due to the uniform BLI scale applied across all images, low tumor signals may not be visually apparent; successful implantation was confirmed relative to baseline BLI, and mice without confirmed tumor implantation were excluded. **B)** Changes in mice body weight determined weekly.

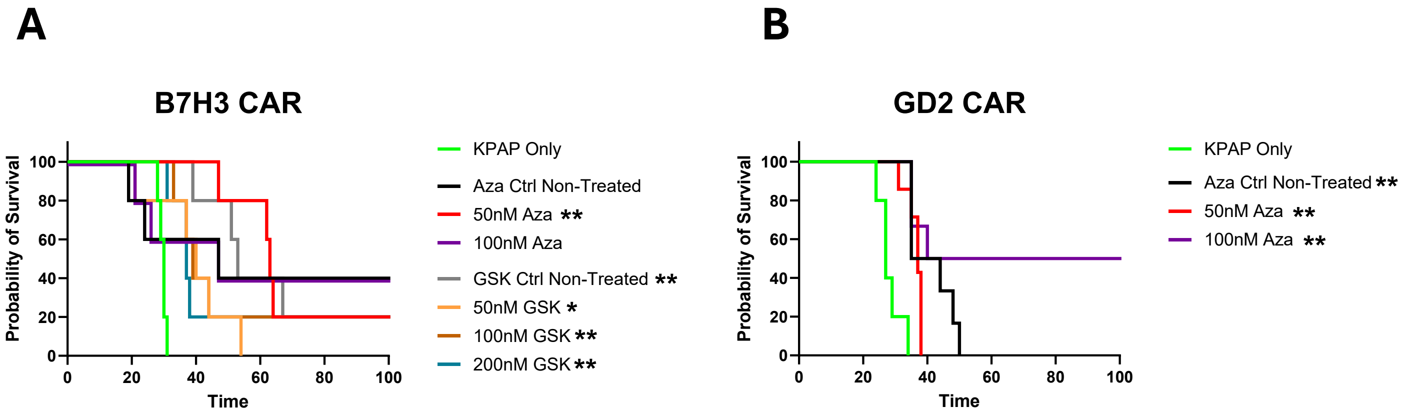


**Supplemental Figure 10. KPAP survival curves with the inclusion of tumor only control. A)** Kaplan-Meier survival curve (n = 5, log-rank Mantel-Cox test with Holm-Šídák correction for multiple comparisons. Asterisks denote significant comparisons versus tumor-only control, *P < 0.05, **P < 0.01. **B)** Kaplan-Meier survival curve (n = 5-7, log-rank Mantel-Cox test with Holm-Šídák correction for multiple comparisons. Asterisks denote significant comparisons versus tumor-only control, **P < 0.01. The ID8 tumor-only group (n=5) represents previously published data from Gomez et al. (2022) and is shown as a reference for typical survival in this model.

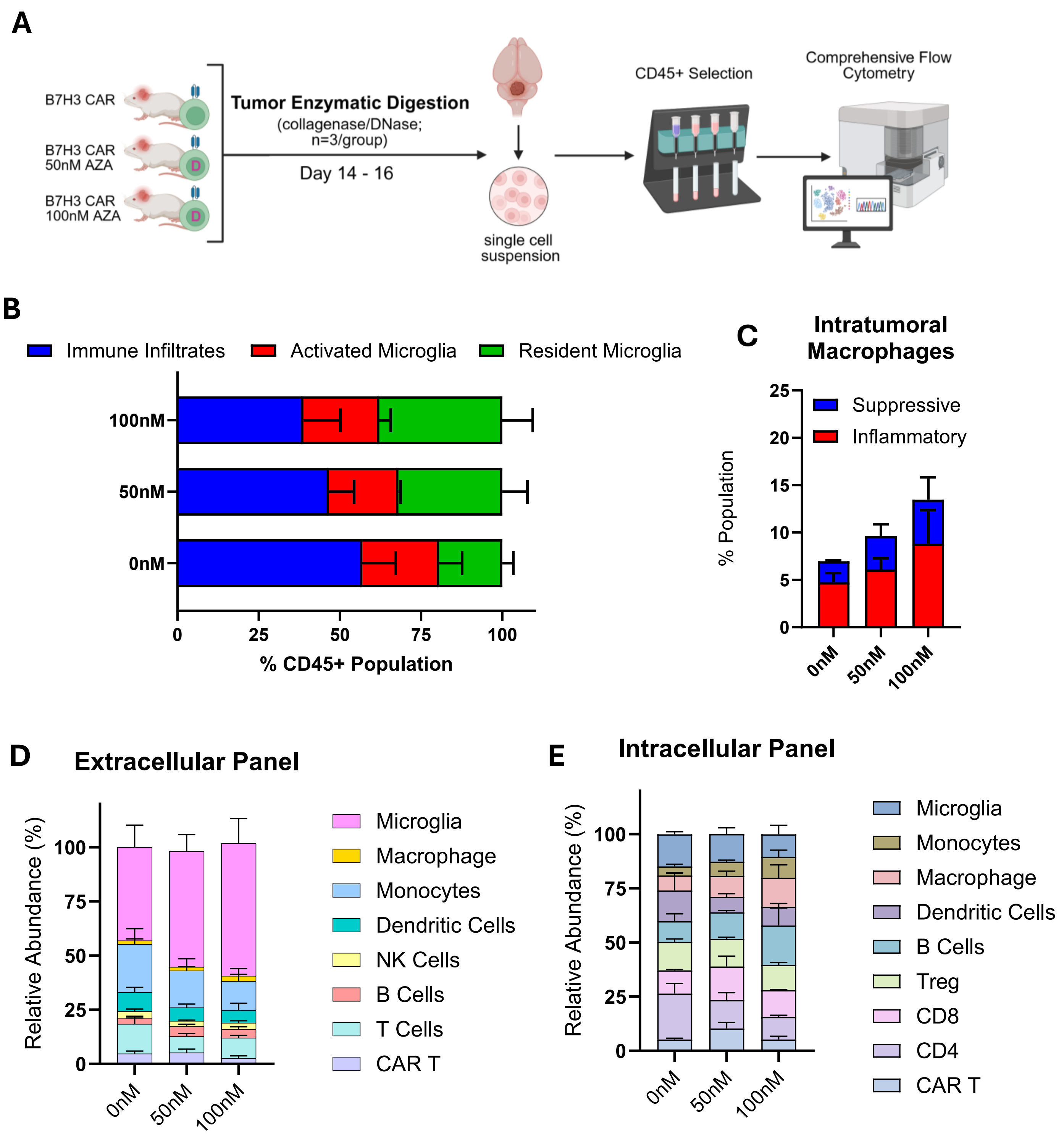


**Supplemental Figure 11. Flow cytometric analysis of the KPAP tumor microenvironment 4 days post CAR T-cell treatment. A)** Experimental scheme. Albino C57BL/6 mice were orthotopically implanted with 1 × 10^5 KPAP cells, followed 10 days later by intratumoral injection of 2 × 10^6 B7H3 CAR T cells primed with 5-Aza. Tumors were collected 4 days after CAR T-cell treatment, dissociated into single-cell suspensions, and analyzed by spectral flow cytometry following CD45^+ enrichment. **B)** Quantification of major myeloid populations, including resident microglia ($CD45^{low}$), activated microglia ($CD45^{low}$ $F4/80^{low}$ $MHCII^{high}$ $CD11b^{low}$), and infiltrating immune cells ($CD45^{high}$). **C)** Quantification of intratumoral macrophage populations within CD45^+ cells using the intracellular staining panel, including suppressive ($ARG^{high}$ $IL\text{-}10^{high}$) and inflammatory ($TNF\alpha^{high}$ $IL\text{-}6^{high}$) populations. **D-E)** Frequencies of major immune cell populations within each treatment group identified using **D)** cell-surface and **E)** intracellular staining panels (n = 3). Immune cell populations were manually annotated based on lineage and phenotypic marker expression summarized in Supplementary Table 4.

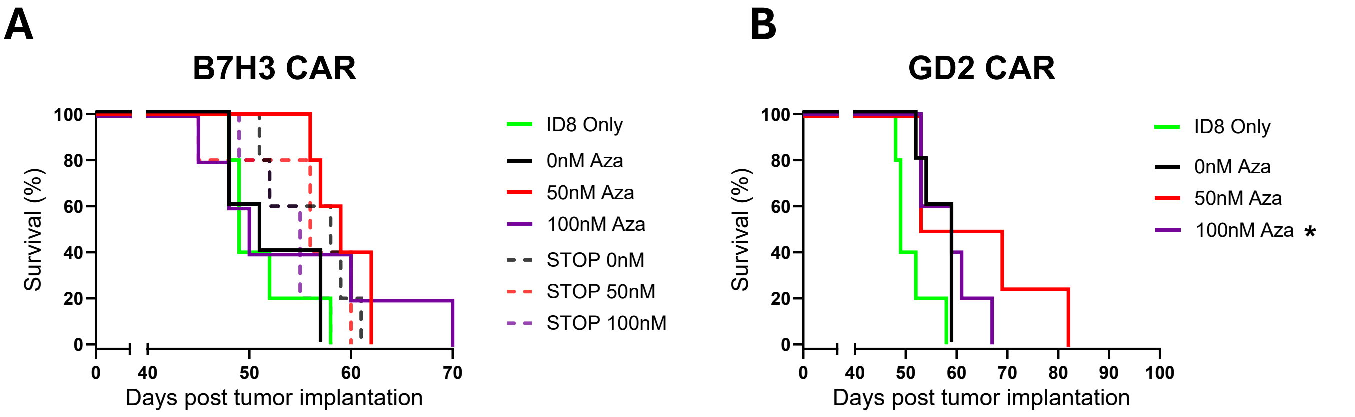


**Supplemental Figure 12. ID8 survival curves with the inclusion of tumor only and STOP controls. A)** Kaplan-Meier survival curve (n = 5, log-rank Mantel-Cox test with Holm-Šídák correction for multiple comparisons. Asterisks denote significant comparisons versus tumor-only control. **B)** Kaplan-Meier survival curve (n = 4-5, log-rank Mantel-Cox test with Holm-Šídák correction for multiple comparisons. Asterisks denote significant comparisons versus tumor-only control, **P < 0.01.

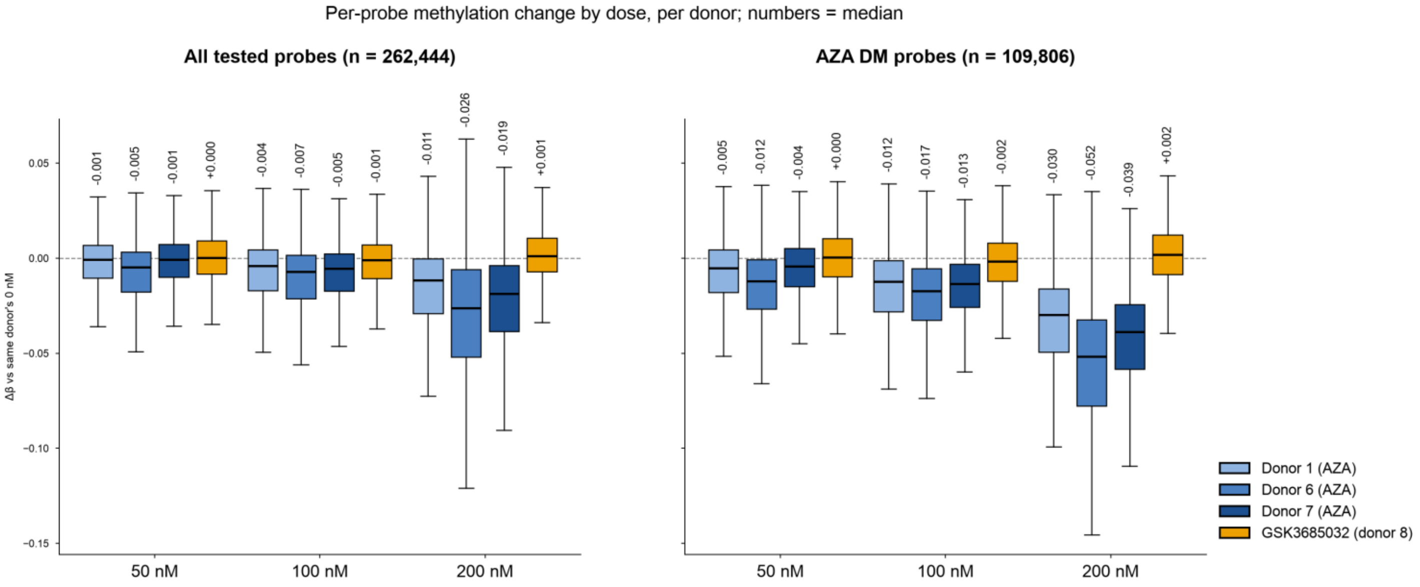


**Supplemental Figure 13. Comparison of Aza- and GSK3685032-associated DNA methylation changes. A)** Methylation changes across all tested probes (n = 262,444) and **B)** Aza differentially methylated (DM) probes (n = 109,806) in individual Aza-treated donors and the GSK3685032-treated donor. Probe-level methylation change (Δβ) at 50, 100, and 200 nM was calculated relative to the corresponding 0 nM condition from the same donor. Boxplots show the distribution of Δβ values, with numbers above each box indicating the median.

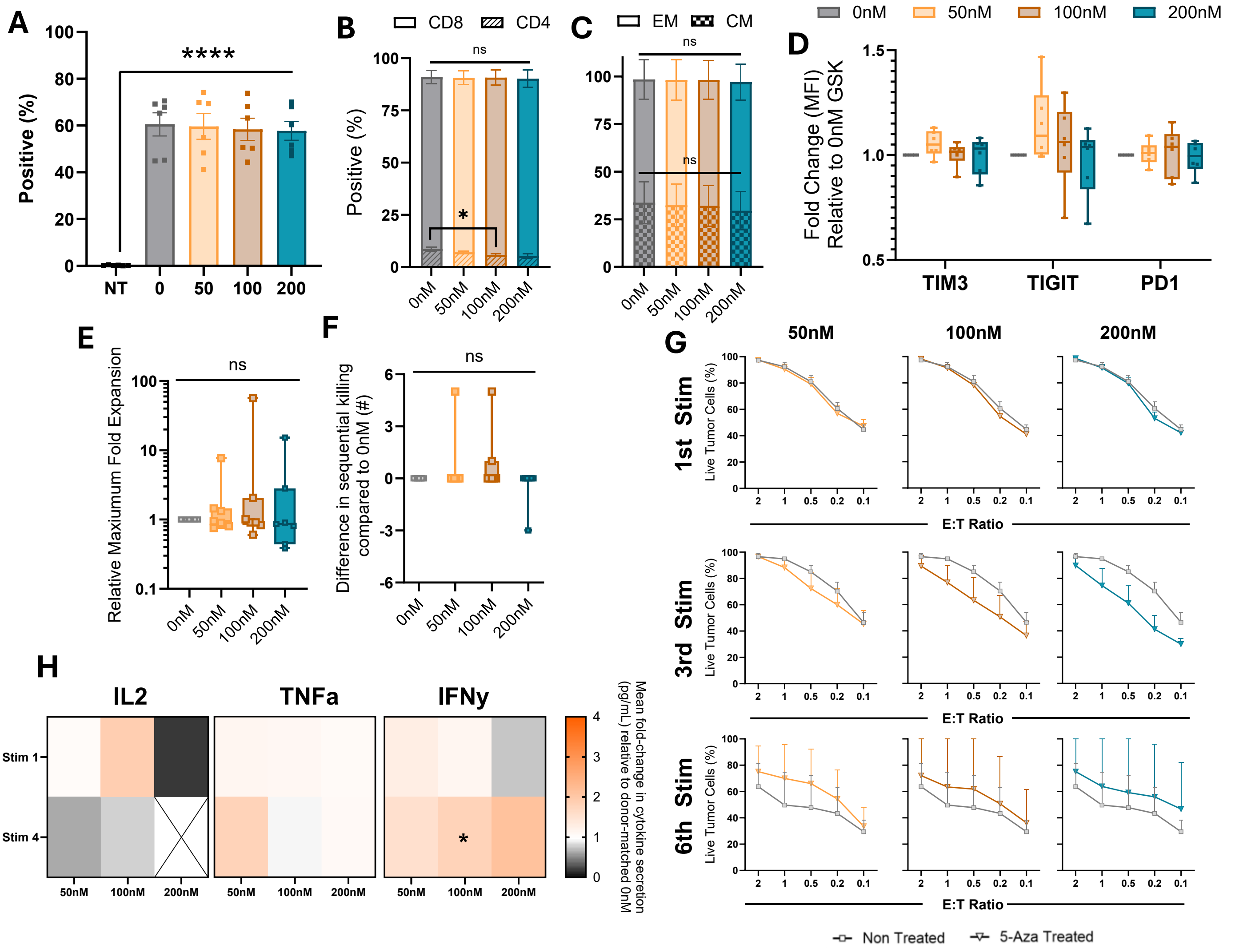


**Supplemental Figure 14. Functional characterization of GSK3685032 treated B7H3 CAR T cells with GL261. A)** Summary plot of %F(ab')2-positive T cells generated in the presence of GSK (n = 6, mean ± SEM, 1-way ANOVA with Tukey's test for multiple comparisons, NT = Non-transduced, *P < 0.05. **B-C)** At day 6 post-transduction, CAR T cells were analyzed by flow cytometry for phenotypic and exhaustion markers. **B)** CD4 and CD8 composition and **C)** effector memory (EM; CD44+/CD62L−) and central memory (CM; CD44+/CD62L+) phenotype distribution of the CD8+ population. Data are shown as mean ± SEM (n = 7) and were analyzed by two-way ANOVA with Tukey's multiple-comparisons test comparing treatment conditions within each phenotype, *P < 0.05. **D)** Fold change in exhaustion marker median fluorescence intensity (MFI) at each 5-Aza concentration, normalized to the donor-matched non-treated control (0 nM). Data are shown as minimum-to-maximum range (n = 7) and were analyzed by one-sample Wilcoxon signed-rank test against a hypothetical value of 1.0. **E)** Maximum fold expansion during co-culture at each GSK concentration relative to the donor-matched untreated (0 nM) control, calculated as the treated maximum divided by the 0 nM maximum (n = 7, minimum-to-maximum range, one-sample Wilcoxon signed-rank test versus a hypothetical value of 1.0). **F)** Difference in the maximum number of sequential GL261 tumor-cell stimulations supported while maintaining CAR T-cell cytotoxic activity relative to the donor-matched untreated (0 nM) control (n = 7, minimum-to-maximum range, one-sample Wilcoxon signed-rank test versus a hypothetical value of 0). **G)** MTS cytotoxicity assay against GL261 tumor cells across E:T ratios following the first (n = 7), third (n = 4) and sixth (n = 3) stimulation rounds. Data are shown as mean ± SEM. **H)** IL-2, TNFa and IFNγ secretion measured by ELISA over consecutive stimulation rounds. Heatmap values represent the median fold change in cytokine secretion relative to the donor-matched untreated (0 nM) control (n = 3, one-sample t test versus a hypothetical value of 1.0, *P < 0.05).

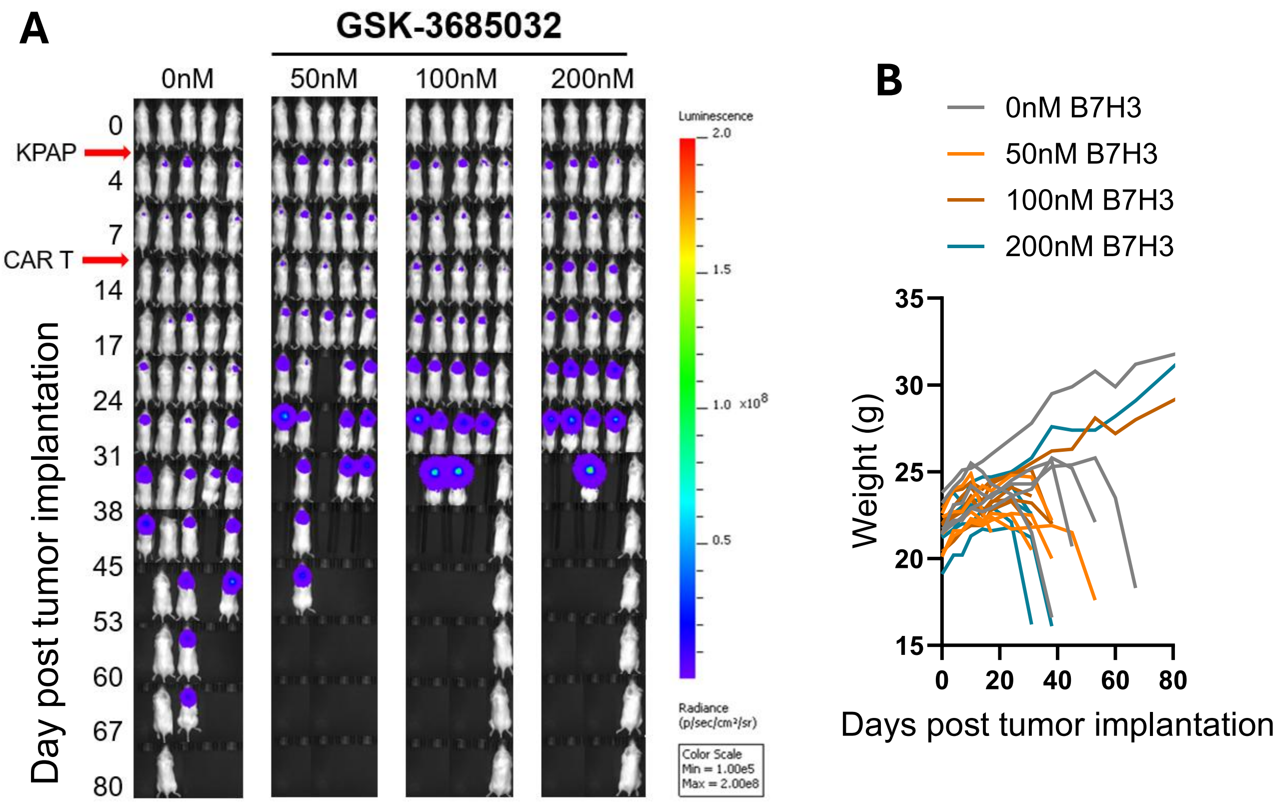


**Supplemental Figure 15. Longitudinal bioluminescent imaging and weight monitoring following GSK-primed B7H3 CAR T cell therapy in the KPAP model.** On day 0, mice were stereotactically implanted with 1 × $10^5$ KPAP tumor cells. On day 10, mice were treated with 1 × $10^6$ B7-H3 CAR T cells primed with GSK-3685032 during manufacturing. **A)** BLI images showing tumor burden at the indicated time points following KPAP implantation and B7-H3 CAR T-cell treatment. Imaging parameters, shown at left, were kept consistent throughout the study. Due to the uniform BLI scale applied across all images, low tumor signals may not be visually apparent; successful implantation was confirmed relative to baseline BLI, and mice without confirmed tumor implantation were excluded. **B)** Changes in mice body weight determined weekly.

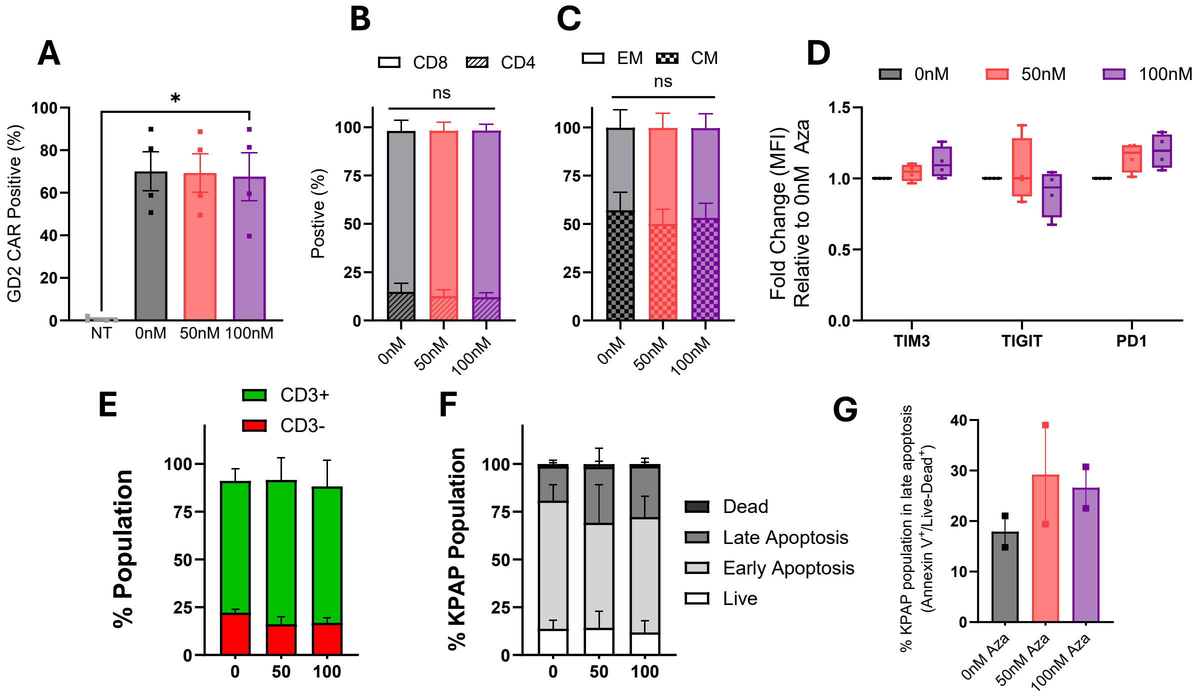


**Supplemental Figure 16. Assessment of GD2 5-Aza treated CAR T cell product. A)** Summary plot of %1A7-positive T cells generated in the presence of 5-Aza (n = 4, mean ± SEM, 1-way ANOVA with Tukey's test for multiple comparisons, NT = Non-transduced, *P < 0.05. **B-C)** At day 6 post-transduction, CAR T cells were analyzed by flow cytometry for phenotypic and exhaustion markers. **B)** CD4 and CD8 composition and **C)** effector memory (EM; CD44+/CD62L−) and central memory (CM; CD44+/CD62L+) phenotype distribution of the CD8+ population. Data are shown as mean ± SEM (n = 4) and were analyzed by two-way ANOVA with Tukey's multiple-comparisons test comparing treatment conditions within each phenotype. **D)** Fold change in exhaustion marker median fluorescence intensity (MFI) at each 5-Aza concentration, normalized to the donor-matched non-treated control (0 nM). Data are shown as minimum-to-maximum range (n = 4) and were analyzed by one-sample Wilcoxon signed-rank test against a hypothetical value of 1.0. **E)** The entire collected co-culture was analyzed by flow cytometry to quantify the relative frequencies of CD3+ CAR T cells and CD3− KPAP tumor cells. Stacked bar plots show the percentage of each population following co-culture with GD2 CAR T-cells primed with 0–100 nM 5-Aza after one stimulation (n =2). **F)** CD3- Tumor cells were classified as live, early apoptotic, late apoptotic, or dead based on Annexin V and live/dead staining. Stacked bar plots show the distribution of tumor cell states following co-culture with B7-H3 CAR T cells primed with 0–100 nM 5-Aza after one stimulation (n =2). **G)** Summary bar plot only showing CD3- KPAP cells in late apoptosis (Annexin V+/Live/Dead+, n = 2).

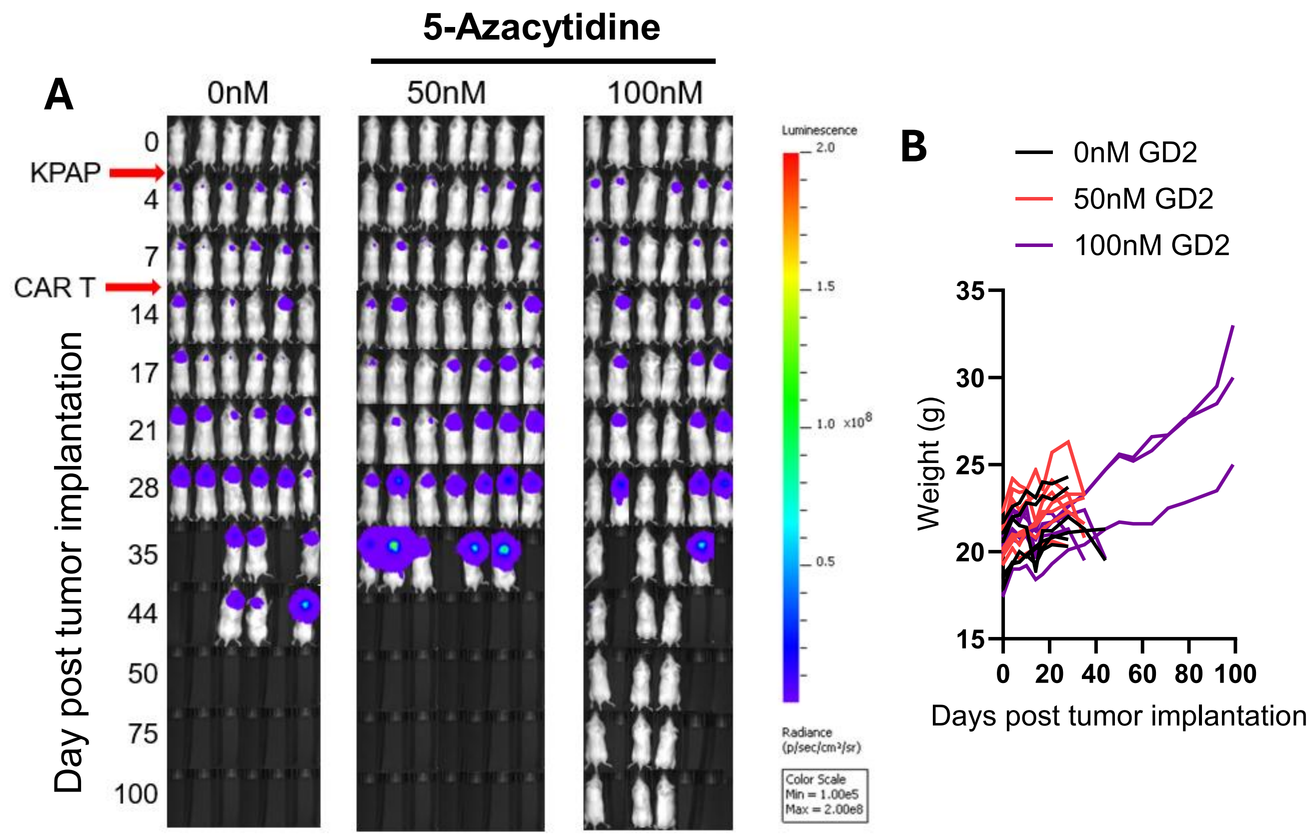


**Supplemental Figure 17. Longitudinal bioluminescent imaging and weight monitoring following 5-Aza-primed GD2 CAR T cell therapy in the KPAP model**. On day 0, mice were stereotactically implanted with 1 × $10^5$ KPAP tumor cells. On day 10, mice were treated with 2 × $10^6$ GD2 CAR T cells primed with 5-Aza during manufacturing. **A)** BLI images showing tumor burden at the indicated time points following KPAP implantation and GD2 CAR T-cell treatment. Imaging parameters, shown at left, were kept consistent throughout the study. Due to the uniform BLI scale applied across all images, low tumor signals may not be visually apparent; successful implantation was confirmed relative to baseline BLI, and mice without confirmed tumor implantation were excluded. **B)** Changes in mice body weight determined weekly.

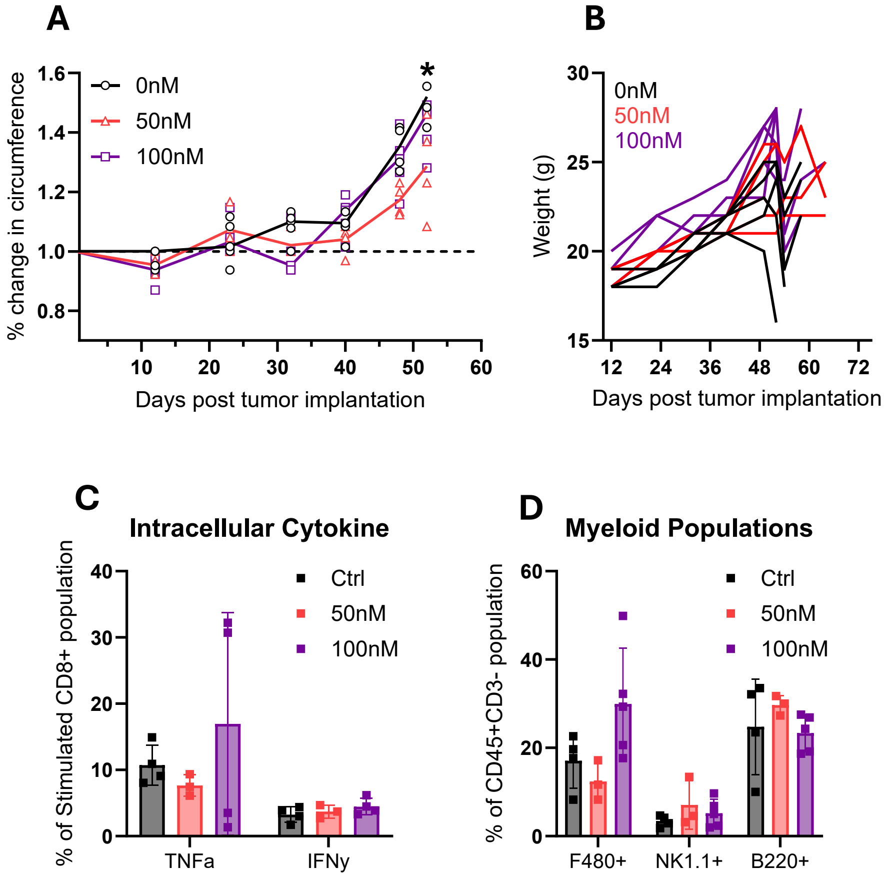


**Supplemental Figure 18. Longitudinal development of ovarian cancer and immune profiling of drained ascites following treatment with 5-Aza primed GD2 CAR T cells. A**) Percent change in abdominal circumference following ID8 Trp53^-/-^ tumor injection. Symbols represent individual mice within each treatment group, the dotted line indicates baseline circumference, and data are presented as mean ± SEM. Statistical significance was assessed using a Kruskal-Wallis test, with the asterisk indicating a significant difference between the untreated (0 nM) and 50 nM 5-Aza-primed GD2 CAR T-cell groups, *P < 0.05. **B)** Weekly changes in mouse body weight. **C)** Quantification of TNFα and IFNγ expression in ascites-derived CD8+ T cells following ex vivo stimulation (n = 3). **D)** Quantification of myeloid and immune cell populations within the CD3−CD45+ fraction of ascites, including macrophages (F4/80+), NK cells (NK1.1+), and B cells (B220+) (n = 3).
