## Supplemental Methods for "DNA Methyltransferase Inhibition During CAR T-cell Manufacturing Enhances Antitumor Activity in Immunocompetent Brain and Ovarian Cancer Models"

### Cell Lines

GL261 murine glioma tumor cells were procured from the Leibniz Institute DSMZ-German Collection of Microorganisms and Cell Cultures. MBR525-1 and KPAP cells were kindly provided by our collaborators Zoe Weaver Ohler (NIH) and Fred Wu (Virginia Tech), respectively. GL261 tumor cells were cultured in complete RPMI media (Cytiva, Marlborough, MA, USA) with 10% fetal bovine serum (FBS), 1% glutamax, 1% penicillin/streptomycin, 1% sodium pyruvate, 1% MEM non-essential amino acids, and 0.1% of 2-mercaptoethanol. MBR525-1 were maintained in complete DMEM (GenClone, Madison, WI, USA) with 10% fetal bovine serum (FBS), 1% penicillin/streptomycin, 2.5% HEPES and 0.2% primocin. KPAP cells were cultured in Neurobasal-A medium (Life Technologies, Carlsbad, CA, USA) supplemented with B-27 (2%), N-2 (1%), GlutaMAX (1%), penicillin-streptomycin (1%), heparin (2 μg/mL), human bFGF (20 ng/mL), and human EGF (20 ng/mL). ID8 murine cell lines (*Trp53*$^{-/-}$) were gifted from Dr Iain McNeish[44] and were cultured in DMEM media (Gibco, Thermo Fisher Scientific, Waltham, MA, USA) with 4% FBS, 1% penicillin/streptomycin, ITS (5 μg/mL insulin, 5 μg/mL transferrin, and 5 μg/mL sodium selenite). Murine T-cells were cultured in complete RPMI media with 10% FBS, 1% glutamax, 1% penicillin/streptomycin, 1% MEM non-essential amino acids, and 0.1% of 2-mercaptoethanol.

### DNMT Inhibitor Preparation

Lyophilized 5-azacytidine (5-Aza; (5-Aza; A2385, Sigma-Aldrich) and GSK3685032 (GSK; E1046, Selleck Chemicals) were reconstituted in dimethyl sulfoxide (DMSO) to generate 500μM stock solutions. To preserve compound stability, stock solutions were aliquoted, stored at −80°C, and discarded following a single freeze-thaw cycle. Prior to use, stock solutions were diluted 1:10 in complete T-cell culture medium to generate 50μM working solutions. Working solutions were added directly to cell culture to achieve final concentrations of 50, 100, or 200nM. The corresponding final DMSO concentrations were 0.01%, 0.02%, and 0.04%, respectively. Untreated control cultures received an equivalent volume of complete T-cell culture medium without drug.

### Western Blotting

CAR T-cells were collected on day 6 post-transduction, washed once with phosphate-buffered saline (PBS), pelleted, and immediately stored at −80°C until analysis. Cell pellets were thawed and lysed in RIPA buffer (Pierce, 89900) supplemented with 1× protease and phosphatase inhibitor cocktail (Pierce, A32961). Lysates were sonicated in an ice-cold water bath sonicator for 8 minutes (8 cycles of 30s on and 30 s off) and clarified by centrifugation. Protein concentration was determined using a Pierce BCA Protein Assay Kit (Thermo Fisher Scientific, 23225) according to the manufacturer's instructions. Equal amounts of protein (15μg per sample) were mixed with NuPAGE™ LDS Sample Buffer (4×; Invitrogen, NP0007) and NuPAGE™ Sample Reducing Agent (10×; Invitrogen, NP0009), denatured at 95°C for 5 minutes, and loaded onto 4-20% Mini-PROTEAN TGX gels (Bio-Rad, 4561093). Proteins were transferred to low-fluorescence PVDF membranes (Bio-Rad, 2023954), which were subsequently blocked with Intercept® (PBS) Blocking Buffer (LI-COR Biosciences, 927-70001). Membranes were incubated with primary antibodies against DNMT1 (Sigma-Aldrich, D4692; 1:250), DNMT3A (Abcam, ab227823; 1:1000), and β-actin (ACTB; Sigma-Aldrich, A5441; 1:3000), followed by IRDye® 680RD Goat Anti-Rabbit IgG (LI-COR Biosciences, 926-68071; 1:5000) and IRDye® 800CW Goat Anti-Mouse IgG (LI-COR Biosciences, 926-32210; 1:5000) secondary antibodies. β-actin was used as a loading control. Fluorescent signal was detected using a ChemiDoc™ MP Imaging System (Bio-Rad, Hercules, CA, USA). Band intensities were quantified using ImageJ software and normalized to β-actin expression.

**Syngeneic Tumor Models**

For orthotopic intracranial models, anesthetized mice were secured in a stereotactic frame. For cortical implantation, a 1-mm burr hole was created and $1\times10^5$ GL261 cells in 2 µL were injected 1 mm anterior and 2 mm right of bregma and 4 mm ventral from the skull surface. Fourteen days later, mice received $1–2\times10^6$ CAR T-cells in 1–2 µL RPMI intratumorally using the same coordinates. Cell dose was adjusted for CAR transduction efficiency. For pontine implantation, $1\times10^5$ KPAP.ffLuc cells in 1 µL Neurobasal-A medium were injected 0.8 mm posterior and 1 mm right of lambda and 5.5 mm ventral from the skull surface. Ten days later, mice received $1–2\times10^6$ CAR T-cells in 1 µL RPMI using the same coordinates.

For ovarian cancer studies, female C57BL/6J mice received $5\times10^6$ ID8 $Trp53^{-/-}$ cells intraperitoneally in 0.5 mL PBS, followed 21 days later by $3\times10^6$ CAR T-cells intraperitoneally. Body weight and abdominal circumference were monitored weekly, and ascites was drained when either increased ≥30% from baseline. Under isoflurane anesthesia, ascites was removed using a 21-gauge needle, followed by up to 5 mL sterile saline subcutaneously. Mice were euthanized after three drainage procedures or earlier for signs of distress, including labored breathing, hypoactivity, or malnutrition.

**Animal Study Design and Reporting**

Mice were maintained under standard institutional housing conditions on a 12-hour light/dark cycle with ad libitum access to food and water. Animals were provided environmental enrichment in accordance with institutional animal care procedures. Animals were acclimated for one week before initiation of experimentation. A total of ~120 mice were used across all in vivo studies. Group-specific sample sizes and exclusions are reported in the corresponding figures and legends. The individual mouse was considered the experimental unit for all *in vivo* studies. Control groups included tumor-only mice and mice receiving untreated CAR T-cell products. Tumor-only controls were shared across studies to minimize unnecessary animal use. Sample sizes were based on experimental feasibility; no formal *a priori* power calculation was performed. For bioluminescent models, mice were allocated and randomized using Microsoft Excel to balance baseline mean flux across treatment groups. For models in which tumor burden could not be directly visualized, mice were randomly allocated without knowledge of tumor burden. Treatment order was alternated across groups to minimize order-related confounding. Investigators were not blinded to treatment allocation or outcome assessment. Animals were excluded only for failure to confirm tumor establishment or for unexpected death occurring before the anticipated tumor-related endpoint and considered unrelated to tumor progression or the experimental intervention. Experiment-specific exclusions and reasons for exclusion are reported in the corresponding figure legends.

**Bioluminescence (IVIS) imaging**

Mice were injected intraperitoneally with AkaLumine hydrochloride (MedChemExpress, Monmouth Junction, NJ) (5 mM in PBS; 100 µL per mouse) 5 minutes before imaging, anesthetized with isoflurane (2.5% delivered in 100% $O_2$ at 0.5 L/min), and imaged with a IVIS Imaging System Series

III (Perkin Elmer, Shelton, CT). The photons emitted from the luciferase-expressing cells were quantified using Living Image software (Caliper Life Sciences, Waltham, MA). Mice were imaged by IVIS prior to tumor implantation to establish baseline signal and on days 4 and 7 post-implantation to monitor tumor establishment. Mice were subsequently imaged weekly, including one day prior to CAR T-cell administration to assess pretreatment tumor burden. Following CAR T-cell treatment, imaging was performed on days 4 and 7 and weekly thereafter to evaluate treatment response and long-term tumor control.

Flow Cytometry

**Murine T-cell Flow Cytometry-Based Characterization**
Murine CAR T-cell phenotype and transduction efficiency were evaluated by flow cytometry. Samples were acquired on a CytoFLEX flow cytometer (Beckman Coulter, Brea, CA, USA) and analyzed using FlowJo software (BD Biosciences, Ashland, OR, USA). Viable cells were identified using LIVE/DEAD™ Fixable Aqua Dead Cell Stain (Thermo Fisher Scientific, Cat. No. L34966). B7H3 CAR expression was detected using Alexa Fluor® 647 AffiniPure $F(ab')_2$ Fragment Goat Anti-Human IgG, $F(ab')_2$ fragment specific (Jackson ImmunoResearch, Cat. No. 109-606-006), while GD2 CAR expression was detected using the anti-idiotype monoclonal antibody 1A7 (Frederick National Laboratory for Cancer Research, Part No. 50475) followed by APC-conjugated anti-mouse IgG1 secondary antibody (BioLegend, Cat. No. 406609). Non-transduced T-cells were used as negative controls for CAR detection. Phenotypic characterization included staining for CD4 (BD Pharmingen, Cat. No. 552051), CD8α (BD Pharmingen, Cat. No. 553030), CD44 (BD Pharmingen, Cat. No. 553134), CD62L (BD Pharmingen, Cat. No. 560513), TIM-3 (BioLegend, Cat. No. 134019), TIGIT (BioLegend, Cat. No. 142106), and PD-1 (BioLegend, Cat. No. 135220). Data were analyzed through sequential gating to assess viability, CAR expression, T-cell lineage, differentiation status, and expression of exhaustion-associated markers.

**Tumor Antigen Profiling**
Utilized Tumor cell lines were evaluated for surface expression of B7H3 and GD2 by flow cytometry. B7H3 expression was assessed using anti-CD276 (B7H3) Alexa Fluor® 647 antibody (BD Pharmingen, Cat. No. 562862) with Rat IgG2a, κ Alexa Fluor® 647 isotype control (BD Pharmingen, Cat. No. 557690). GD2 expression was assessed using GD2 Alexa Fluor® 647 antibody (BioLegend, Cat. No. 357318) with Mouse IgG2a, κ Alexa Fluor® 647 isotype control (BioLegend, Cat. No. 400234).

**Annexin assay**
To assess CAR T-cell-mediated cytotoxicity against non-adherent tumor cells, KPAP cells were seeded in low-attachment 24-well plates at $5x10^5$ cells/well and allowed to form neurospheres for 2-4 hours. CAR T-cells were added at $1x10^6$ cells/well (2:1 effector-to-target [E:T] ratio) in a total volume of 2 mL and co-cultured for 72 hours. Cells were then harvested, counted and stained with anti-CD3 (BD Biosciences, Cat. No. 562600) for 30 minutes on ice and then stained with near-IR Live/Dead viability dye (Invitrogen, Cat. No. L34976) for 20 minutes on ice. Cells were then resuspended in 1X binding buffer (Invitrogen, Cat. No. V13246) and stained with fluorochrome-conjugated Annexin V (Invitrogen, Cat. No. A35110) for 15 minutes at room temperature. After staining, additional binding buffer was added to samples and data was acquired on a CytoFLEX flow cytometer (Beckman

Coulter, Brea, CA, USA) within 1 hour of staining. Data was analyzed using FlowJo (BD Biosciences, Ashland, OR, USA). The proportions of CD3+ and CD3− cells were quantified, and tumor-cell apoptosis and death were assessed within the CD3− population. CAR T-cell-only and KPAP-only controls were used to establish gating. When KPAP neurosphere elimination was evident, $1x10^6$ of the collected CAR T-cells were restimulated with fresh KPAP cells. After an additional 72 hours of co-culture, Annexin V analysis was repeated, and serial restimulation was continued until CAR T-cell cytotoxic activity was no longer evident.

**Immune Profiling of Ovarian Cancer Ascites**

Ascites was centrifuged at 500×g for 8 minutes at 4°C, and erythrocytes were removed by ACK lysis. Cells were washed, counted, and plated at $1x10^6$ cells/mL in RPMI containing 10% FBS and 1X penicillin-streptomycin. Cells were either treated with eBioscience Protein Transport Inhibitor Cocktail (Thermo Fisher Scientific, Cat. No. 00-4980-03) or Cell Stimulation Cocktail with protein transport inhibitors (Thermo Fisher Scientific, Cat. No. 00-4975) at a final concentration of 1X and incubated for 4-6 hours prior to flow cytometry staining. Cells were resuspended in FACS buffer (PBS + 1% FBS) and treated with Brilliant Stain Buffer (BD Biosciences, Cat. No. 563794), True-Stain Monocyte Blocker (BioLegend, Cat. No. 426103), and TruStain FcX anti-mouse CD16/32 antibody (BioLegend, Cat. No. 101320). Surface staining was then performed for 40–60 minutes at 4°C with a lymphoid panel; anti-CD3e-AF700 (BioLegend, Cat. No. 152316), anti-CD45-PerCP/Cy5.5 (BioLegend, Cat. No. 103132), anti-CD90.1-BV750 (BD Biosciences, Cat. No. 746928), anti-CD4-BV650 (BioLegend, Cat. No. 100555), anti-CD8α-APC/Fire750 (BioLegend, Cat. No. 100766), anti-CD62L-BV421 (BioLegend, Cat. No. 104436), anti-CD44-PE/Cy7 (BioLegend, Cat. No. 103029), and anti-NK1.1-BV605 (BioLegend, Cat. No. 108753), or a myeloid panel; anti-CD11b-BV785 (BioLegend, Cat. No. 101243), anti-CD11c-BV421 (BioLegend, Cat. No. 117343), anti-F4/80-PE (BioLegend, Cat. No. 123110), anti-I-A/I-E-FITC (BioLegend, Cat. No. 107606), anti-CD80-BV650 (BioLegend, Cat. No. 104732), and anti-CD40-PE/Cy7 (BioLegend, Cat. No. 124622). Cells were fixed using the FoxP3/Transcription Factor Staining Buffer Set (Thermo Fisher Scientific, Cat. No. 00-5523-00) for 1 hour on ice and subsequently stained intracellularly with anti-IFN-γ-FITC (BioLegend, Cat. No. 505806), anti-TNF-α-PE/Dazzle 594 (BioLegend, Cat. No. 506346), and anti-FoxP3-AF647 (BD Biosciences, Cat. No. 563486) for 40–60 minutes at 4°C. Samples were acquired on a Cytek Aurora spectral flow cytometer and analyzed using FlowJo v10 to quantify cell populations within the ascites.

**Immune Profiling of KPAP Tumor Burden**

To evaluate CAR T-cell persistence and immune populations within the TME, tumor-bearing brains were harvested at 4 days post CAR T-cell treatment. Pontine tissue containing tumor and adjacent normal brain was minced and enzymatically digested with collagenase (1 mg/mL) and DNase I (50 U/mL) for 60 minutes at 37 °C. Digests were mechanically dissociated and sequentially filtered through 70-m and 40-m strainers to generate single-cell suspensions, followed by ACK-mediated erythrocyte lysis and PBS washes. Additional filtration and low-speed centrifugation were applied as needed to reduce cellular debris. CD45+ immune cells were enriched by magnetic selection using mouse CD45 MicroBeads (Miltenyi Biotec, 130-097-153). CD45+ cells were the resuspended at 1e6 cells per mL in 5mL FACS tubes and prepared for spectral flow cytometry. CD45+ cell samples were assessed with both a 35-Color and 21-Color panel and antibodies utilized are detailed in Table 1a and b. The panels were validated on a Cytek Aurora 5 Laser (UV-V-B-YG-R) using primary cells isolated from murine brain tissue to replicate experimental conditions. Single color controls were generated from both cells and beads, providing a unique unmixing profile of cells and beads for both panels (Table 1a and b). Unmixing was confirmed using full stained samples that were used for the single-color control generation.

**Sample Staining: 35-Color Panel**

Cells were washed in 2mL of PBS and centrifuged at 400xg for 5 minutes. Supernatant was decanted and cells were resuspended in a 1:150 working solution of 200uL UV Blue viability dye and incubated for 30 minutes on ice. After incubation, cells were washed in 2mL of MACS quant running buffer at 400xg for 5 minutes. Supernatant was decanted and cells were resuspended in 100uL of MACS quant buffer with 5uL of True Stain FCR Block and incubated for 30 minutes in the dark on ice. While cells were incubating in FCR block, the 36-color surface master mix was made with antibodies and dilutions detailed in Table 2. After incubation, cells were stained directly with 104uL of the master mix and incubated for 30 minutes in the dark on ice. Cells were then washed with 2mL of MACS quant buffer at 400xg for 5 minutes, resuspended in 500uL of MACS quant and collected on the Cytek Aurora.

**Sample Staining: 21-Color Panel**

Prepared FACS samples were incubated at 37°C for 4 hours with 2uL/mL of PMA/Ionomycin (BioLegend, Cat No. 423301) to allow for accumulation of intracellular cytokines. After incubation, cells were washed with 2mL of PBS and stained with UV Blue viability dye and FCR block as previously described. An extracellular master mix was made with antibodies and dilutions detailed in Table 3a. Cells were stained with 69uL of the surface mix and incubated for 30 minutes in the dark on ice. Cells were then washed with 2mL of MACs quant running buffer at 400xg for 5 minutes and the supernatant discarded. Cells were resuspended in 1mL of 1x Fix Concentrate (Biolegend Cat#73158), incubated at room temperature in the dark for 45 minutes, then washed twice with 2mL of True Nuclear 1x Perm Buffer (Biolegend Cat#73162) at 400xg for 5 minutes. The intracellular master mix was made according to the dilutions in Table 3b, and cells were stained with 68uL of the intracellular mix. Cells were incubated at room temperature in the dark for 30 minutes, washed twice with 2mL of True Nuclear 1x Perm Buffer at 400xg for 5 minutes, resuspended in 500uL of Perm Buffer and collected on the Cytek Aurora.

**OMIQ Analysis**

Samples were collected following QC and calibration of the Aurora and assessment of unstained samples. Once the samples were collected, the unmixing profile was applied and the data exported as unmixed FCS files. These files were then uploaded into OMIQ.ai for further analysis. Once in OMIQ, samples were evaluated by gating for Cells/Singlets/Live/Dead/CD45+ cells. Once CD45+ cells were identified, subsampling was performed to eliminate all debris and non CD45+ cells. This population of cells was analyzed using T-distributed Stochastic Neighbor Embedding (tSNE) and clustering to investigate subsets of CD45+ cells and compare relative abundance across samples. Following clustering, immune cell populations were manually annotated based on lineage and phenotypic marker expression, with cluster identities summarized in Table 4. For phenotypic analysis of infiltrating CD90.1+ CAR T-cells, normalized median fluorescence intensity (MFI) values for each marker were exported from OMIQ and subsequently standardized by z-score normalization across samples.

| Cells | Beads |
|---|---|
| CD45 BUV395 | CD24 BUV496 |
| Live Dead Blue | CD44 BUV563 |
| CD11c BUV615 | CD11b BUV661 |
| CD62L eFluor450 | CD103 BUV805 |
| CD3 BV480 | CD163 BV421 |
| CD8 Spark Violet 538 | CD80 BV510 |
| Ly6C BV570 | CD206 BV711 |
| CD86 BV605 | CD90.1 BV750 |
| CD64ab BV650 | CD276 PE-Dazzle 594 |
| XCR1 BV786 | MHCII PE-Fire 810 |
| CD19 Spark Blue 550 | CD123 APC |
| CD4 Spark Blue 574 | CD69 Spark NIR 685 |
| Ly6G PerCP | |
| PD-1 BB700 | |
| CD25 PerCP-eFluor710 | |
| CD209a PE | |
| F4/80 Spark YG 593 | |
| CD127 PE-Cy5 | |
| CD49b PE-Cy5.5 | |
| TIM-3 PE-Cy7 | |
| CTLA-4 APC-R700 | |
| LAG-3 APC-Fire 750 | |
| NK1.1 APC-Fire 810 | |

Table 1a. 35 color unmixing profile.

| Cells | Beads |
|---|---|
| Live Dead Blue | 1L-17A BUV395 |
| CD11c BUV615 | CD45 BUV496 |
| CD3 BUV661 | IFNγ BUV737 |
| CD4 BUV805 | RORγt BV421 |
| CD11b BV570 | CD25 BV510 |
| TCRβ BV605 | TNFα BV650 |
| F4/80 BV711 | CD90.1 BV750 |
| CD8 BV786 | IL6 PE |
| FOXP3 PerCP | 1L10 PE-Cy7 |
| AHR PE-CF594 | ARG1 APC |
| B220 AF700 | |

Table 1b. 21 color unmixing profile.

| Marker | Color | Vendor | Catalog # | Vol. per 100uL | Dilution |
|---|---|---|---|---|---|
| CD45 | BUV395 | BD | 564279 | 0.5 | 1:200 |
| CD80 | BV510 | Biolegend | 104741 | 1 | 1:100 |
| CD4 | Spark Blue 574 | Biolegend | 100489 | 2 | 1:50 |
| MHC II | PE Fire 810 | Biolegend | 107667 | 0.1 | 1:1200 |
| CD8a | Spark Violet | Biolegend | 155019 | 1 | 1:100 |
| Ly6G | PerCP | Biolegend | 127653 | 1 | 1:100 |
| CD123 | APC | Invitrogen | 17-1231-82 | 0.625 | 1:160 |
| CD25 | PerCP eF710 | Invitrogen | 46-0251-82 | 2 | 1:50 |
| CD69 | Spark NIR 685 | Biolegend | 104557 | 2 | 1:50 |
| CD86 | BV605 | BD | 563055 | 2 | 1:50 |
| CD209a | PE | Biolegend | 833003 | 2 | 1:50 |
| LAG3 | APC Fire 750 | Biolegend | 125239 | 1 | 1:100 |
| CD11b | BUV661 | BD | 612977 | 1 | 1:100 |
| F4/80 | Spark YG 593 | Biolegend | 157311 | 1 | 1:100 |
| NK1.1 | APC Fire 810 | Biolegend | 156519 | 1 | 1:100 |
| CD206 | BV711 | Biolegend | 141727 | 2 | 1:50 |
| CD276 | PE Dazzle 594 | Biolegend | 135611 | 2 | 1:50 |
| CD163 | BV421 | Biolegend | 155309 | 2 | 1:50 |
| CD127 | PECy5 | Biolegend | 135015 | 2 | 1:50 |
| CD24 | BUV496 | BD | 612953 | 0.2 | 1:500 |
| CD62L | eF450 | Invitrogen | 48-0621-80 | 1 | 1:100 |
| XCR1 | BV785 | Biolegend | 148225 | 3.3 | 1:30 |
| CD49b | PECy5.5 | Invitrogen | 35-5971-80 | 1 | 1:100 |
| CD103 | BUV805 | BD | 741948 | 2 | 1:50 |
| CD19 | Spark Blue 550 | Biolegend | 115565 | 1 | 1:100 |
| TIM3 | PECy7 | Biolegend | 119715 | 2 | 1:50 |
| CTLA-4 | APC R700 | BD | 565778 | 1 | 1:100 |
| CD3 | BV480 | BD | 741562 | 5 | 1:20 |
| PD-1 | BB700 | BD | 566515 | 2 | 1:50 |
| CD44 | BUV563 | BD | 741227 | 0.33 | 1:300 |
| CD11c | BUV615 | BD | 751265 | 1 | 1:100 |
| CD64a/b | BV650 | BD | 740622 | 2 | 1:50 |
| CD90.1 | BV750 | BD | 746961 | 0.4 | 1:250 |
| Viability | UV Blue | Invitrogen | L23105 | 0.66 | 1:150 |
| Ly6C | BV570 | Biolegend | 128030 | 1 | 1:100 |

Table 2. 35 color surface panel and master mix dilutions.

| Marker | Color | Vendor | Catalog # | Vol. for 100uL | Dilution |
|---|---|---|---|---|---|
| F4/80 | BV711 | BD | 565612 | 2 | 1:50 |
| CD25 | BV510 | Biolegend | 102041 | 1.7 | 1:60 |
| TCRb | BV605 | BD | 562840 | 1 | 1:100 |
| B220 (CD45R) | AF700 | Invitrogen | 56-0452-80 | 1 | 1:100 |
| CD11b | BV570 | Biolegend | 101233 | 0.5 | 1:200 |
| CD8 | BV786 | BD | 563332 | 1 | 1:100 |
| CD3 | BUV661 | BD | 741562 | 5 | 1:200 |
| CD45 | BUV496 | BD | 749889 | 0.33 | 1:300 |
| CD90.1 | BV750 | BD | 746961 | 0.4 | 1:250 |
| CD4 | BUV805 | BD | 612900 | 0.5 | 1:200 |
| CD11c | BUV615 | BD | 751265 | 1 | 1:100 |
| Viability | UV Blue | Invitrogen | L23105 | 0.66 | 1:150 |
| Monocyte Blocking Buffer | | | | 5 | 1:20 |

Table 3a. 21 color surface panel and master mix dilutions.

| Marker | Color | Vendor | Catalog # | Vol. for 100uL | Dilution |
|---|---|---|---|---|---|
| IL-10 | PECy7 | Biolegend | 505025 | 2 | 1:50 |
| IL-6 | PE | Biolegend | 504503 | 2 | 1:50 |
| AHR | PE-CF594 | BD | 565790 | 2 | 1:50 |
| IFNg | BUV737 | BD | 612769 | 2 | 1:50 |
| RORgt | BV421 | BD | 562894 | 2 | 1:50 |
| TNFa | BV650 | Biolegend | 506333 | 2 | 1:50 |
| ILI7A | BUV395 | BD | 565246 | 2 | 1:50 |
| FoxP3 | PerCPCy5.5 | Invitrogen | 45-5773-82 | 2 | 1:50 |
| ARG1 | APC | Invitrogen | 17-3697-80 | 2 | 1:50 |
| Brilliant Buffer | | | | 50 | 1:2 |

Table 3b. 21 color intracellular panel and master mix dilutions.

| Immune Lineage | Cell Subset | Flow Markers |
|---|---|---|
| Microglia | Resident | CD45 (low/-) F4/80(-) CD64(Low/-) |
| | Activated | CD45(Low) F4/80(Low/-) CD64 (+) Ly6C(+) MHCII(+) |
| Macrophage | Supressive | F4/80 (+) ARG (+) IL10 (+) CD163 (+) CD206 (+) |
| | Inflammatory | F4/80 (+) TNFa (+) IL6 (+) CD11b (+) CD64 (+) MHCII (+) CD80 (+) |
| Lymphoid Cells | NK Cells | CD3 (-) NK1.1 (+) |
| | B lymphocytes | CD3 (-) CD220 (+) |
| T Lymphocytes | CD4 | CD3 (+) CD4 (+) |
| | CD8 | CD3 (+) CD8 (+) |
| | Treg | CD3 (+) CD4 (+) FOXP3 (+) IL17A (+) |
| | CAR T | CD3 (+) CD90.1 (+) |
| Monocytes | | CD11b (+) F4/80 (low) Ly6C (+) Ly6G (-) |
| Dendritic Cells | | CD11c (+) MHCII (+) |

Table 4. Marker-based identification of major immune cell populations within CD45+ cells

Bioinformatic Analysis

The RNA-seq cohort comprised five donors (NS: 18 samples, as two donors lacked the 200nM condition; S1: 12 samples from three donors × four doses). The methylation cohort comprised 24 arrays from four donors; the primary analysis used the 12 AZA-treated B7H3 CAR samples (three donors × four doses). Only one donor was shared between the two cohorts, so RNA-seq and methylation data were integrated at gene- and pathway-level dose-response effects rather than by pairing samples.

**Bulk RNA sequencing**

Total RNA was isolated with the QIAGEN RNeasy Mini Kit. Total RNA integrity was determined using Agilent Bioanalyzer. Library preparation was performed with 100ng of total RNA. Libraries were generated with Watchmaker Library Prep Kit with Polaris Depletion (Watchmaker). Briefly, ribosomal RNA was removed by an RNaseH method and purified with RNAClean beads (Beckman). mRNA was then fragmented in buffer by heating depending on RNA quality per protocol. mRNA was reverse transcribed to yield strand specific cDNA. A second strand and A Tailing reaction was performed to yield fragments with an A base added to the 3’ ends. Illumina sequencing adapters were ligated to the ends. Ligated fragments were then amplified 12 cycles using primers incorporating unique dual index tags. Fragments were sequenced on an Illumina NovaSeq X Plus using paired end reads extending 150 bases and targeting 30M read pairs.

**RNA-seq data processing**

Reads were processed with a Snakemake[1] (v8.20.6) workflow. Read quality was assessed with FastQC[2] (v0.12.1), and adapter and low-quality sequences were removed with Trim Galore[3,4] (v0.6.10; cutadapt v5.2) in paired-end mode. For quality control, trimmed reads were aligned to the GRCm39 primary assembly with GENCODE[5] vM32 annotation using STAR[6] (v2.7.0e; --*outFilterMultimapNmax 100 --winAnchorMultimapNmax 200*), sorted with samtools[7] (v1.21), and summarized with Picard[8] CollectRnaSeqMetrics (v3.2.0) and MultiQC[9] (v1.25.2).
Gene-level counts were generated with SQuIRE[10] (v0.9.9.92) and run from a container image via Apptainer (v1.3.0). The SQuIRE index was built from the GRCm39 primary assembly and GENCODE vM32 annotation (sjdbOverhang 150), and the UCSC mm39 RepeatMasker annotation[11] was used for the Clean step. SQuIRE Map aligned trimmed reads with STAR (v2.5.3a; read length 151), and

SQuIRE Count quantified gene expression with StringTie[12] (v1.3.3) using reverse-stranded library settings. Per-sample gene counts were merged into a single gene-by-sample count matrix.

**Differential expression analysis**

Differential expression was performed in R (v4.5.3; Bioconductor 3.22) with DESeq2[13] (v1.50.2). Non-stimulated (NS) and Stimulated (S1) samples were analysed separately; the NS analysis was the primary analysis. Genes with at least 10 counts in at least 3 samples were retained. AZA dose was modelled as a continuous covariate (dose in nM divided by 100) with donor as a blocking factor (design ~ donor + dose), so that the tested coefficient is the log2 fold change per 100nM AZA. Log2 fold changes were shrunk with apeglm[14] (v1.32.0), and P values were adjusted with the Benjamini–Hochberg (BH) method. Genes with adjusted $P < 0.05$ and an absolute shrunken log2 fold change > 0.5 were considered differentially expressed (DE). Ensembl identifiers were mapped to gene symbols and Entrez identifiers with org.Mm.eg.db[15] (v3.22.0).

**RNA-seq gene set enrichment analysis**

Gene set enrichment analysis (GSEA) of the NS dose-response model was performed with clusterProfiler[16] (v4.18.4) *gseGO*, which uses fgsea[17] (v1.36.2). Genes were ranked by the DESeq2 Wald statistic for the dose coefficient. All genes with a Wald statistic were included. GO Biological Process[18] (GO:BP) terms with 15–500 genes were tested, P values were BH-adjusted, and terms with FDR < 0.05 were considered significant. Redundant terms were reduced for display with clusterProfiler *simplify* (semantic-similarity cutoff 0.7, Wang measure; GOSemSim[19] v2.36.0).

**DNA methylation arrays**

Genomic DNA was isolated from treated cells with the Zymo Research Quick-DNA MiniPrep Plus Kit. 500ng of genomic DNA was bisulfate converted using the Zymo EZ 96 DNA Methylation kit. Denatured genomic DNA sample was incubated with Illumina whole-genome amplification (WGA) master mix RA1 at 37°C overnight in an isothermal reaction. After the fragmentation reaction, the purification of the DNA was performed with the blue precipitation reagent and isopropanol and the plate was centrifuged at 4°C for 20 minutes to pellet the DNA, followed immediately by a decanting operation: firmly invert the plate onto an absorbent pad to remove the supernatant and air-dry the visible blue pellets on a tube rack for 1 hour. The DNA pellets were then incubated with a hybridization buffer in the oven at 48°C for 1 hour to fully resuspend the DNA sample. The GDA BeadChip Hybridization was done overnight at 48°C. A single-base enzymatic extension was performed using labeled nucleotides for the hybridized DNAs on the BeadChip, followed by chip scanning in the iScan System. The scanned data was loaded into the GenomeStudio software (v1.0) for chip quality assessment. The chip QC was confirmed without any warning signs. The methylation data was exported for downstream genotyping analysis.

**Methylation array preprocessing**

IDAT files from all 24 arrays were processed with sesame[21] (v1.28.1; sesameData v1.28.0). Signal intensities were read with *readIDATpair* and preprocessed with *prepSesame* (prep code QCDPBM) to remove technical artifacts. Probe quality masking was restricted to the reference-issue and design-issue masks, so that probes in repetitive sequence were retained. We applied dye-bias correction, background detection masking with pOOBAH ($P < 0.05$), noob (normal-exponential out-of-band) background correction to correct for background fluorescence, and Infinium I/II design matching. Beta values were then obtained with *openSesame* (*func* = *getBetas*). No between-array normalization or batch corrections were applied, because donor and BeadChip were confounded. Sample quality and outliers were assessed by principal component analysis of the 20,000 most variable CpG probes. In total, 277,292 probes were tested.

Probe coordinates from the sesameData MM285 manifest (mm10) were converted to mm39 with the UCSC[11] mm10ToMm39 chain file using rtracklayer[22] (v1.70.1) *liftOver*, keeping probes that mapped one-to-one, at the same width, to standard chromosomes (288,653 of 288,655 probes, including all tested probes). Gene annotation used the same GENCODE vM32 models as the RNA-seq analysis, with a TxDb built with txdbmaker (v1.6.2).

**Differential methylation analysis**

The primary analysis used the 12 AZA-treated B7H3 CAR samples (three donors × four doses). Probe-level dose responses were modelled on M-values with limma[23] (v3.66.0; *lmFit* and *eBayes*) using the design ~ donor + dose, with AZA dose as a continuous covariate. Differentially methylated regions (DMRs) were identified with DMRcate[24] (v3.6.0) from a manually constructed CpG-annotated object, as MM285 is not supported by *cpg.annotate* (*dmrcate* parameters: lambda = 1000, C = 2, consec = FALSE, pcutoff = "fdr", min.cpgs = 2). Effect sizes were expressed on the beta scale as the fitted change from 0 to 200nM AZA, and DMRs with an absolute mean beta difference > 0.10 were retained (507 DMRs). Because DMRcate annotates genes only for mm10, DMR gene overlaps were reassigned from the GENCODE vM32 gene models.

**Methylation gene set enrichment analysis**

Gene set enrichment of DMRs was tested with dmGsea[25] (v1.0.2) *gsProbe*. The significant probes were the tested probes inside the 507 DMRs (1,055 probes) and the background was all tested probes that mapped to a gene. Probes were assigned to genes if they lay between 1.5 kb upstream of the transcription start site and the gene end (promoter and gene body), and gene identifiers were converted to Entrez with org.Mm.eg.db. Enrichment was tested with Fisher's noncentral hypergeometric distribution (BiasedUrn v2.0.12), which weights each gene's probability of selection by its number of probes. GO:BP terms with 15–500 genes were tested and terms with FDR < 0.05 were considered significant. dmGsea v1.0.2 computes the upper-tail probability as $P(X > k)$ instead of $P(X \geq k)$; this was corrected by replacing the internal function *p.gs* before analysis. Threshold- and rank-based dmGsea modes were not used because AZA induced widespread hypomethylation (38% of probes at FDR < 0.05, 99.7% of them hypomethylated), which saturated those tests.

**Gene-level methylation values**

For gene-level integration, each gene was assigned the probes in its promoter and gene body (the dmGsea mapping described above). The gene-level methylation value was the median over these probes, so that long genes with many probes were not favoured. A gene was classified as differentially methylated if at least one of its probes lay inside one of the 507 DMRs (443 genes).

**Integration of RNA-seq and methylation data**

Overlap between DE genes and differentially methylated genes was tested with a one-sided Fisher's exact test (SciPy[26] v1.17.1) among genes measured on both platforms. Overlap between enriched pathways was tested in the same way among GO:BP terms tested in both the RNA-seq GSEA and the methylation dmGsea analysis, with significance defined as FDR < 0.05 in each analysis. The

GO:BP terms significant in both analyses were clustered by Wang semantic similarity (GOSemSim) using hierarchical clustering (ward.D linkage on 1 − similarity) and cut into five clusters. For each shared term, the number of DE genes annotated to the term or its descendant terms and the GSEA leading-edge size were reported.

For heatmaps of DE genes, counts were transformed with the DESeq2 variance-stabilizing transformation (blind = TRUE), and donor effects were removed with limma *removeBatchEffect* (batch = donor, design = ~ dose), which preserves the dose effect. Heatmaps of DMRs showed the mean beta value over each DMR's), donor-adjusted in the same way. For the effector gene set, log2-normalized counts (RNA-seq) and gene-level median beta values (methylation) were donor-adjusted as above and row-scaled. For each candidate gene, the Pearson correlation between the RNA-seq and methylation dose-response trajectories (donor-adjusted means at each of the four doses) was calculated.

**Cytokine gene analysis**

To compare expression between stimulation conditions, DESeq2 size factors were estimated once across all 30 RNA-seq samples (same gene filter), and normalized counts were obtained by dividing raw counts by these size factors. Dose-response statistics were taken from the condition-specific models described above. For methylation, the gene-level median beta value of *Il2*, *Ifng*, *Tnf* and *Ifnb1*, and of the *Ifna* gene family pooled as one unit (*Ifna1–16* and *Ifnab*; the pseudogene *Ifna-ps1* excluded), was modelled by ordinary least squares (beta ~ donor + dose) and reported as the fitted change from 0 to 200nM. To assess these genes genome-wide, the same gene-level model was fitted to every gene with mapped probes, using ordinary and limma-moderated t-statistics with BH correction across genes.

**Comparison of GSK3685032 and azacitidine**

GSK3685032 was tested in a single donor, so these analyses compare that donor's dose response with the AZA response and do not generalize to GSK3685032. The analyses used the 16 B7H3 CAR arrays (three AZA donors and the GSK3685032donor, four doses each). Across the 507 AZA DMRs, the mean beta value over each region's probes (probes without missing values in all 16 samples) was expressed as the change from the same donor's 0nM sample, and a per-donor ordinary least-squares slope on dose was calculated. For an unsupervised comparison, the 500 most variable autosomal CpG probes were selected after subtracting each donor's mean, which removes donor, sex and BeadChip baselines.

The two drugs' dose responses were compared with limma on M-values of autosomal probes without missing values, using the design ~ donor + dose(AZA) + dose(GSK), where each drug's dose is zero in samples treated with the other drug. The contrast dose(GSK) − dose(AZA) was tested (10 residual degrees of freedom), and DMRs were identified with DMRcate using the same parameters and |mean beta difference| > 0.10 filter as the primary analysis.

**Statistical analysis and software**

Unless stated otherwise, P values were adjusted with the BH method and FDR < 0.05 was considered significant. Analyses were run in R (v4.5.3; Bioconductor 3.22) and Python (v3.11). Figures were produced with matplotlib[27] (v3.11.0), matplotlib-venn[28] (v1.1.2), NumPy[29] (v2.4.6), pandas (v3.0.3), ComplexHeatmap[30] (v2.26.1) and circlize[31] (v0.4.18).

**Data and code availability**

RNA-seq FASTQ files and the gene count matrix, and methylation array IDAT files and preprocessed signal and beta values, have been deposited in GEO GSE353449. Code to reproduce all analyses and figures from raw data, together with exact software environments, is available on our lab GitHub (https://github.com/Chiappinelli-Lab/AZA_CAR-T_methylation_rnaseq.git). Two original preprocessing steps were not deterministic: multithreaded StringTie quantification in SQuIRE, and sesame Infinium I/II design matching, for which no random seed was set. The reported results are therefore reproduced exactly from the deposited count matrix and preprocessed methylation data. A complete rerun from raw data gives the same conclusions (the same 111 DE genes and 26 shared GO:BP terms) with minor numerical differences (for example, 517 rather than 507 DMRs).

2. Babraham Bioinformatics - FastQC A Quality Control tool for High Throughput Sequence Data. https://www.bioinformatics.babraham.ac.uk/projects/fastqc/.

3. FelixKrueger/TrimGalore: Consistent adapter and quality trimming for NGS, with extra functionality for RRBS data. https://github.com/FelixKrueger/TrimGalore.

4. Martin, M. Cutadapt removes adapter sequences from high-throughput sequencing reads. *EMBnet.journal* **17**, 10–12 (2011).

5. Mudge, J. M. *et al.* GENCODE 2025: reference gene annotation for human and mouse. *Nucleic Acids Res.* **53**, D966–D975 (2025).

6. Dobin, A. *et al.* STAR: ultrafast universal RNA-seq aligner. *Bioinformatics* **29**, 15–21 (2013).

7. Li, H. *et al.* The Sequence Alignment/Map format and SAMtools. *Bioinformatics* **25**, 2078–2079 (2009).

8. Picard Tools - By Broad Institute. https://broadinstitute.github.io/picard/.

28. konstantint/matplotlib-venn: Area-weighted venn-diagrams for Python/matplotlib. https://github.com/konstantint/matplotlib-venn/tree/master.

29. Harris, C. R. *et al.* Array programming with NumPy. *Nature* **585**, 357–362 (2020).

30. Gu, Z., Eils, R. & Schlesner, M. Complex heatmaps reveal patterns and correlations in multidimensional genomic data. *Bioinformatics* **32**, 2847–2849 (2016).

31. Gu, Z., Gu, L., Eils, R., Schlesner, M. & Brors, B. circlize implements and enhances circular visualization in R. *Bioinformatics* **30**, 2811–2812 (2014).
